# Molecular basis for protection and cross-protection by human antibodies targeting the parainfluenza virus hemagglutinin-neuraminidase protein

**DOI:** 10.64898/2026.03.03.709347

**Authors:** Katelyn D. McCaffrey, Behrouz Ghazi Esfahani, Mohamed A. Elbehairy, Anna L. McCormick, Jarrod J. Mousa

## Abstract

Human parainfluenza viruses (PIVs) are a leading cause of respiratory illness, particularly in vulnerable populations where infection can lead to severe disease. Despite their clinical impact, there are currently no licensed vaccines or effective antiviral treatments available. PIVs have two large surface proteins, the fusion and hemagglutinin-neuraminidase (HN) proteins, both of which are targets of neutralizing antibodies. In this study, we identified and characterized two human monoclonal antibodies (mAbs), 5217-2 and 5217-9, which bind recombinant PIV3 HN protein, bind PIV3-infected cells, and are neutralizing in vitro. We determined the binding epitopes of the PIV3 HN-specific mAbs via biolayer interferometry and found mAb 5217-9 targets a previously defined neutralizing epitope while mAb 5217-2 binds a unique epitope, enabling a more complete understanding of the antigenic landscape. To further understand the newly defined epitope, we determined a cryo-electron microscopy (cryo-EM) structure of mAb 5217-2, which revealed an epitope adjacent to the PIV3 HN protein active site. We also determined the structure of the previously discovered anti-PIV3 HN mAb PIV3HN-09, which was previously shown to be partially protective in vivo. In a hamster challenge model of PIV3, mAb 5217-2 was determined to significantly reduce lung viral titers, demonstrating its protective capacity. Furthermore, as the site 2-directed mAb PIV3HN-05 was previously shown to cross-neutralize PIV1, we evaluated its protective efficacy in an animal challenge model with PIV1, which demonstrated a reduction in lung viral titers. Overall, these findings provide new insights into the antigenic epitopes on the PIV3 HN protein to support structure-based vaccine design efforts and demonstrate new protective mAbs for both PIV3 and PIV1.

## Introduction

Human parainfluenza viruses (PIVs) are a leading cause of acute respiratory infection (ARI) in children, immunocompromised individuals, and the elderly^1^. PIVs can cause both upper and lower respiratory tract infections, typically presenting with cold-like symptoms, or in more severe cases can result in croup, bronchitis, pneumonia, which can lead to hospitalization^1,2^. PIV infections are often overlooked due to the annual epidemics of respiratory syncytial virus (RSV) and influenza virus; however, PIVs are the second most common cause of ARIs in children after RSV^2,3^. PIVs tend to severely affect young children, with those under five years of age exhibiting the highest PIV-associated hospitalization rates^1,4,5^. Among these cases, a substantial proportion occurs in infants, particularly those under one year of age, contributing significantly to infant morbidity and mortality^3,4^. In elderly and immunocompromised populations, PIV infection frequently progresses from upper to lower respiratory tract disease and is associated with high rates of pneumonia, prolonged viral shedding, and increased mortality^6,7^. These individuals are also particularly susceptible to nosocomial transmission and outbreaks in nursing homes^7,8^.

There are five known serotypes of parainfluenza viruses that infect humans: PIV1, PIV2, PIV3, PIV4a, and PIV4b^1,9^. These serotypes differ in their epidemiology and clinical severity. Among them, PIV3 has been associated with lower respiratory tract illness, with symptoms including bronchiolitis and pneumonia, and has been consistently linked to the highest hospitalization rates, reflecting its potential for causing severe respiratory disease^10–12^. PIV1 follows closely behind and is a major contributor to hospital admissions in children under one year of age, most frequently presenting as croup and cold-like symptoms, but it can progress to more severe disease requiring hospitalization^10,11^. In comparison, PIV2 is generally associated with upper respiratory illness, with common cold-like symptoms and lower rates of hospitalization^12,13^. PIV4a and PIV4b remain understudied due to limited historical testing; however, recent studies suggest that these viruses causes respiratory disease comparable to PIV3, with infection of the lower respiratory tract and associated with severe outcomes^4,14^.

The PIVs are negative-sense, single-stranded RNA viruses belonging to the *Paramyxoviridae* family. PIV1 and PIV3 are classified under the genus *Respirovirus*, while PIV2 and PIV4 fall within the genus R*ubulavirus*^1,13^. The RNA genome of PIVs encodes at least six viral proteins: nucleocapsid protein (NP), phosphoprotein (P), matrix protein (M), fusion glycoprotein (F), hemagglutinin neuraminidase glycoprotein (HN), and RNA polymerase (L)^15,16^. The PIV HN and F surface glycoproteins play critical roles in initiating infection. The PIV HN protein is a type II integral membrane protein composed of a stalk region and a globular head domain, which can assemble as a homodimer or homotetramer^17,18^. Unlike many other receptor-binding glycoproteins in the Paramyxovirus family, PIV HN possesses dual functionality: it uses hemagglutination activity to bind sialic acid on host cells and neuraminidase activity to cleave sialic acid and release viral particles during budding^16,19–21^. In addition to mediating attachment to sialic acid-containing receptors, the PIV HN protein plays a critical role in activating the F protein^22^. The PIV F protein mediates the fusion of the viral envelope with the host cell membrane, enabling viral entry^21–23^. Upon activation, the PIV F protein undergoes a conformational change that exposes its buried fusion peptide, enabling the fusion of the viral and host cell membranes^23,24^. Both the PIV HN and F proteins are major targets for neutralizing antibodies, making them key candidates for therapeutic and vaccine development^19^. The virus relies heavily on the PIV HN protein functions, including receptor binding, receptor cleavage, and PIV F protein activation, making it a strategic target for interventions aimed at disrupting receptor binding, thus preventing membrane fusion. While the PIV F protein has been the focus of many antiviral strategies, the multifunctional nature of the PIV HN protein highlights its potential as a more advantageous therapeutic target^19,25,26^.

Currently, there are no approved vaccines for PIVs, despite decades of research efforts. Multiple vaccine platforms have been evaluated in both preclinical and clinical trials, including live-attenuated, vector-based, subunit, and mRNA vaccine candidates^27^. Among the most promising candidates, rPIV3cp45, a recombinant, live-attenuated PIV3 vaccine, has shown safety in infants and young children and induces an immune response against the virus^27–30^. Other serotypes have also been clinically tested: a live-attenuated PIV1 vaccine candidate, rPIV-1/84/del170/942A, was safe in adults and children but was over-attenuated, resulting in lower immune responses in PIV1-seronegative children^31^. Similar findings were reported for the live-attenuated PIV2 vaccine candidates^32^. More recently, mRNA vaccine platforms have been explored due to the success of the COVID-19 mRNA vaccines. An mRNA-based candidate targeting both PIV3 and human metapneumovirus (hMPV) was shown to induce immune responses against both viruses in adults and seropositive children^33,34^.

Increasing attention is being directed toward monoclonal antibodies (mAbs) as a potential preventative and therapeutic strategy. mAbs are highly specific proteins that can target conserved viral or bacterial antigens and have been reported with favorable safety profiles across numerous clinical studies^35–37^. In the case of PIVs, where immunocompromised individuals are at a heightened risk for severe disease, passive immunization with mAbs offers a safe approach for providing immediate, antigen-specific protection^38^. Significant advances have been made in mAb research for respiratory viruses, such as RSV, which has three commercialized mAbs to reduce disease burden in infants^27^: Palivizumab, nirsevimab, and clesrovimab all target the RSV fusion protein and have been approved to provide passive immunity in high-risk and/or healthy infants^39–41^. In addition, several mAbs targeting PIVs have demonstrated protective efficacy in animal models^27^. PIA174, a mAb targeting the PIV3 F protein, has been shown to reduce viral load in both the lungs and nasal passages of cotton rats^42,43^. Furthermore, the broadly reactive mAb 3×1 targets the F proteins of both PIV3 and PIV1, and when administered as a cocktail with the mAb MxR (which neutralizes RSV and hMPV), has been shown to significantly reduce lung viral titers against all four viruses^44^. The success of mAb-based therapies offers a promising strategy for preventing and treating PIV infections, particularly in populations at high risk of severe disease.

The absence of effective treatments allows human parainfluenza viruses to remain a significant threat to vulnerable populations. In this paper, we identified two new neutralizing human mAbs, 5217-2 and 5217-9, targeting the HN protein of PIV3. Each of these mAbs was found to target a distinct epitope on the PIV3 HN protein, and we were able to characterize the structural features of these epitopes via epitope binning and cryo-electron microscopy (cryo-EM). Prophylactic administration of mAb 5217-2 significantly reduced PIV3 replication in vivo. We also determined the cryo-EM structure of the previously discovered protective mAb PIV3HN-09 in complex with the PIV3HN protein. Finally, we determined that the previously discovered PIV1/PIV3 cross-neutralizing mAb PIV3HN-05 protects against PIV1 in vivo.

## Results

### Binding and neutralizing properties of PIV3HN-targeting mAbs

Peripheral blood mononuclear cells (PBMCs) were isolated from generally healthy human subjects and sorted for PIV3 HN-targeting B cells. Single-cell RNA sequencing of these B cells was performed using the 10x Genomics Chromium platform followed by next-generation sequencing to recover paired heavy and light chain mAb sequences^45–47^. We identified and recombinantly produced two mAbs that were specific to the PIV3 HN protein, termed 5217-2 and 5217-9. The binding of mAbs 5217-2 and 5217-9 to the PIV3 HN protein was confirmed via enzyme-linked immunosorbent assay (ELISA). Both mAbs demonstrated binding to the PIV3 HN protein, with mAb 5217-2 exhibiting an EC_50_ of 17 ng/mL and mAb 5217-9 exhibiting an EC_50_ of 21 ng/mL **(Figure 1A)**. The mAbs were subsequently tested for their ability to neutralize PIV3 *in vitro* using a plaque reduction neutralization assay. This approach quantifies how effectively a mAb can inhibit viral infection by tracking the number of plaques, which are areas of cell death, formed in a cell monolayer. The degree of plaque reduction reflects the neutralizing strength of the mAb. mAb 5217-2 exhibited an IC_50_ of 3 ng/mL against PIV3, while mAb 5217-9 had an IC_50_ of 70 ng/mL, both of which are compared to the previously discovered anti-PIV3 F mAb PIA174 that had an IC_50_ of 15 ng/mL **(Figure 1B)**^43^.

**Figure 1.**
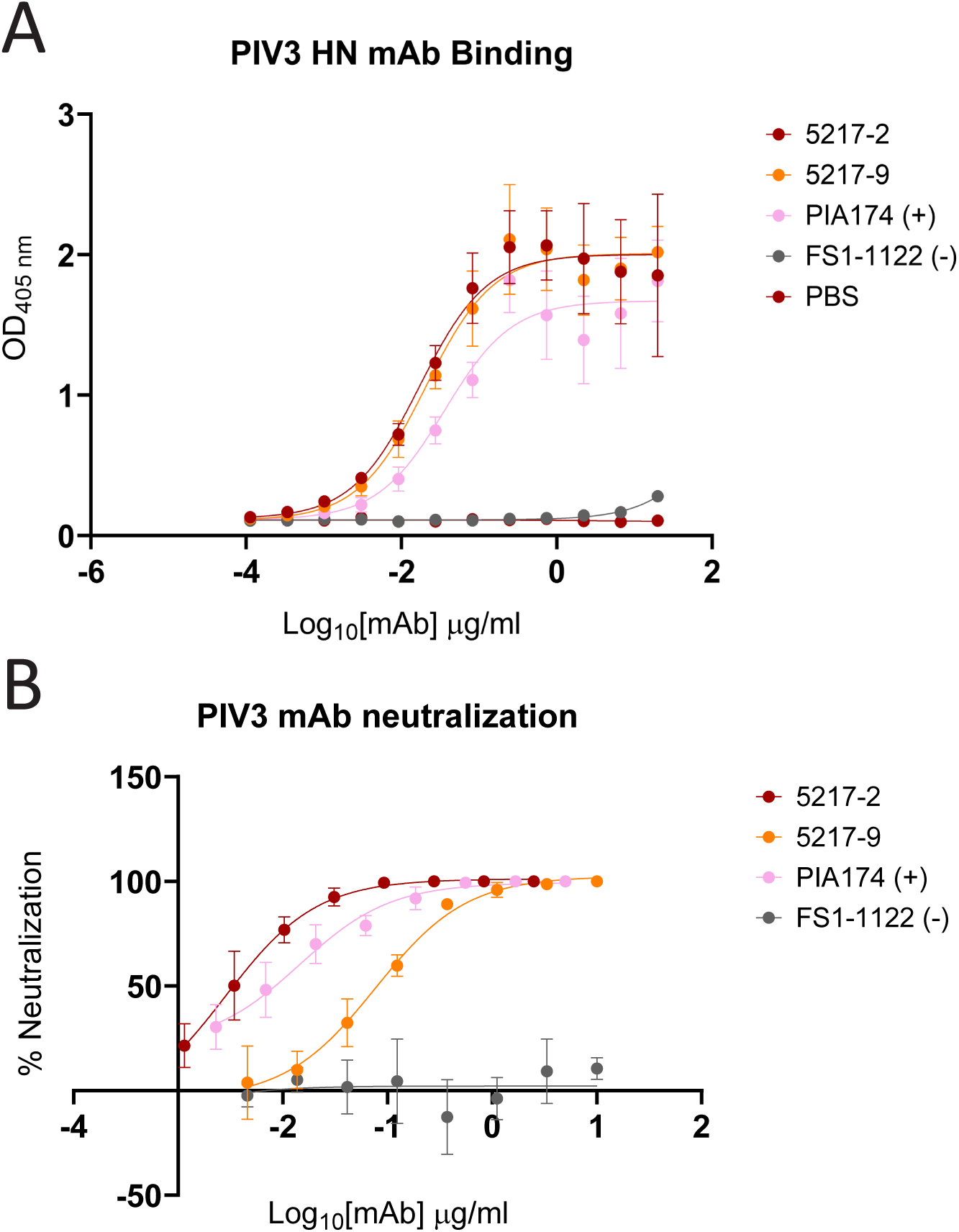
ELISA binding and virus neutralization of isolated mAbs. (A) ELISA binding curves of isolated mAbs, 5217-2 and 5217-9, against the PIV3 HN protein. Optical density at 405 nm (OD_405_) was measured using a microplate reader. Data points are the average of four technical replicates from one experiment and are representative of two biological replicates. The data is presented using mean values +/- standard deviation. (B) Plaque reduction neutralization assay of the two isolated mAbs against PIV3. Data points are the average of three technical replicates from one experiment and are representative of two biological replicates. The data is presented using mean values +/- standard deviation.

To confirm that our mAbs recognize the native full-length PIV3 HN protein, we evaluated their binding to PIV3-infected LLC-MK2 cells via flow cytometry. This technique allows detection of viral proteins as they appear on the surface of infected cells, providing insights into whether the epitopes remain accessible during infection. Both mAbs 5217-2 and 5217-9 showed staining of PIV3-infected cells, comparable to the positive control anti-PIV3 F mAb PIA174. The absence of staining by the anti-pneumococcal mAb PhtD3 confirms the specificity of the interaction **(Figure 2A)**^48^. These data indicate that the epitopes recognized by mAbs 5217-2 and 5217-9 are preserved in the native structure of the PIV3 HN protein and are exposed during infection, supporting the functional relevance of the epitopes targeted by mAbs 5217-2 and 5217-9.

**Figure 2.**
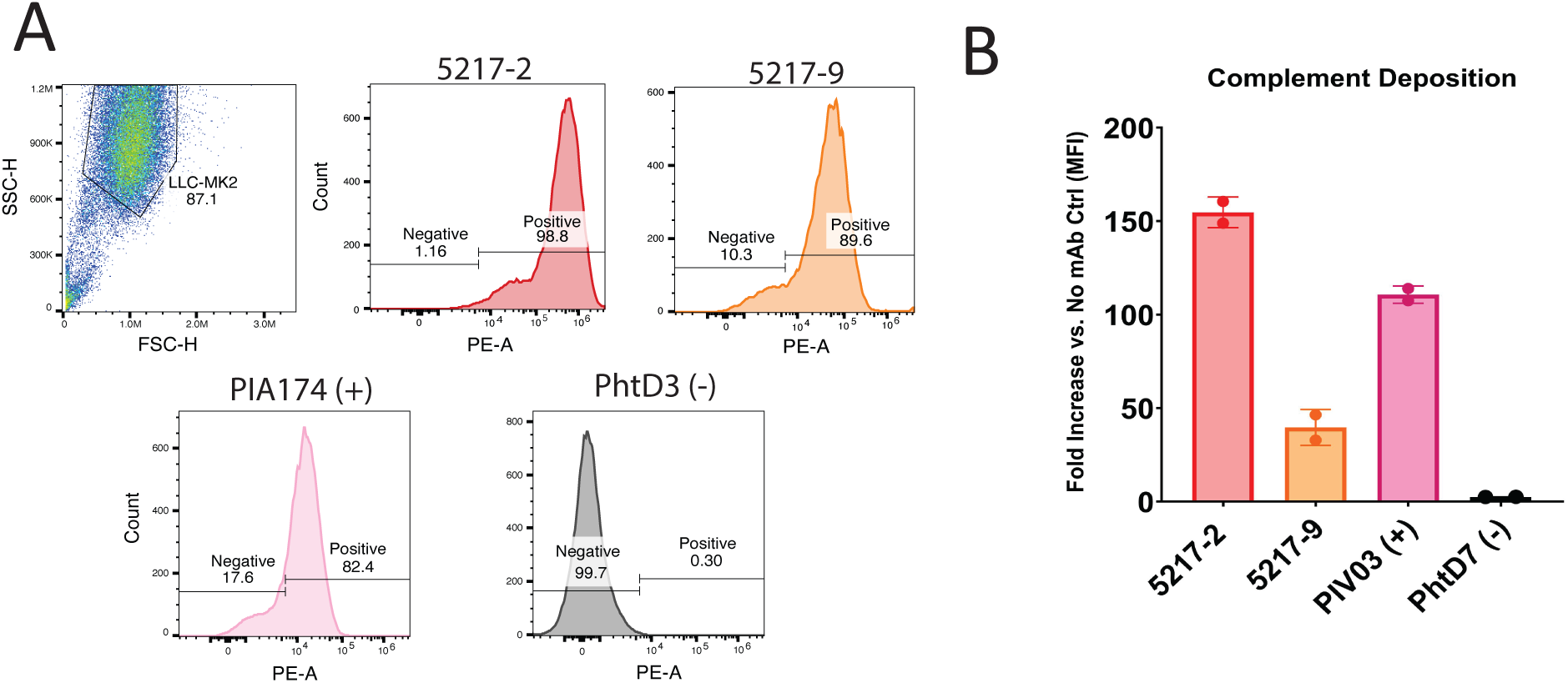
Antibody binding to virally infected cells and associated complement-dependent cytotoxicity assay. (A) LLC-MK2 cells were infected with PIV3 and stained with PE-conjugated PIV3 HN-specific mAbs 5217-2 and 5217-9. The anti-F mAb PIA174 was used as a positive control and a mAb against *Streptococcus pneumoniae*, PhtD3, was used as a negative control. Both the HN-specific mAbs and anti-F mAbs showed binding to PIV3-infected cells. The data represents three technical replicates from one experiment and are representative of two biological replicates. (B) NeutrAvidin beads were coated with biotinylated PIV3 HN protein and incubated with individual mAbs. Guinea pig complement C3 was then added to the immune complexes. Complement deposition was analyzed using the Cytek Aurora flow cytometer and measured relative to C3 deposition in the absence of antibody. Data points are the average of two technical replicates from one experiment and are representative of two biological replicates. The data is presented using mean values +/- standard deviation.

#### Complement deposition mediated by PIV3HN-targeting mAbs

Beyond their ability to recognize and neutralize pathogens through their Fab regions, antibodies also play a crucial role in activating innate immune defenses. Through their Fc region, antibodies can interact with complement proteins, initiating a cascade that leads to the formation of the membrane attack complex (MAC) and ultimately cell lysis of the target cell^49,50^. It was previously shown that anti-PIV3 HN mAbs can initiate this pathway, and Site 2-specific mAbs demonstrated the highest complement deposition activity^46^. To determine whether the newly generated mAbs could initiate this pathway, we evaluated their capacity to promote C3 deposition when introduced to the PIV3 HN antigen. When complement is recruited to the antigen-antibody complex, C3 becomes deposited on the surface and can be detected using a FITC-conjugated anti-C3 antibody. Among the mAbs tested, 5217-2 induced the highest level of complement deposition, resulting in a 150-fold increase in C3 signal relative to the no mAb control. This level of activity exceeded that of the previously described PIV3HN-03, which had shown the strongest complement deposition within its original panel^46^. In contrast, mAb 5217-9 elicited a weaker response of 40-fold increase. As expected, the anti-pneumococcal mAb PhtD7 did not promote C3 deposition **(Figure 2B)**^48^.

#### Epitope mapping reveals a novel binding site on the PIV3 HN protein

To further investigate the antigenic binding profiles of the newly characterized mAbs alongside those previously identified, epitope mapping was performed via competitive biolayer interferometry (BLI), a technique that measures interactions by analyzing interference patterns^51^. By examining changes in the interference pattern, we were able to determine whether antibodies competed for the same binding region when introduced sequentially. The goal was to determine whether the newly isolated mAbs recognized unique epitopes or shared binding sites with the previously described mAbs. The epitope binning experiment was performed in a pairwise combinatorial format, in which each mAb was tested against itself and every other antibody in the panel. Previously, three epitope groups on the PIV3 HN protein, termed antigenic Site 1, Site 2, and Site 3, were mapped using this method^46^. The addition of the newly generated mAbs into the binning assay led to the discovery of a new binding site designated as antigenic Site 4. Notably, mAb 5217-2 demonstrated exclusive binding to this new epitope, as evidenced by its unique binding profile and lack of competition with the Site 1-3 antibodies. This suggests that the PIV3 HN protein possesses an additional accessible binding site that has not been defined. In contrast, mAb 5217-9 was found to compete with mAbs classified within Site 1, such as PIV3HN-13 **(Figure 3)**. Together, these binning results expand our understanding of the antigenic landscape of the PIV3 HN protein.

**Figure 3.**
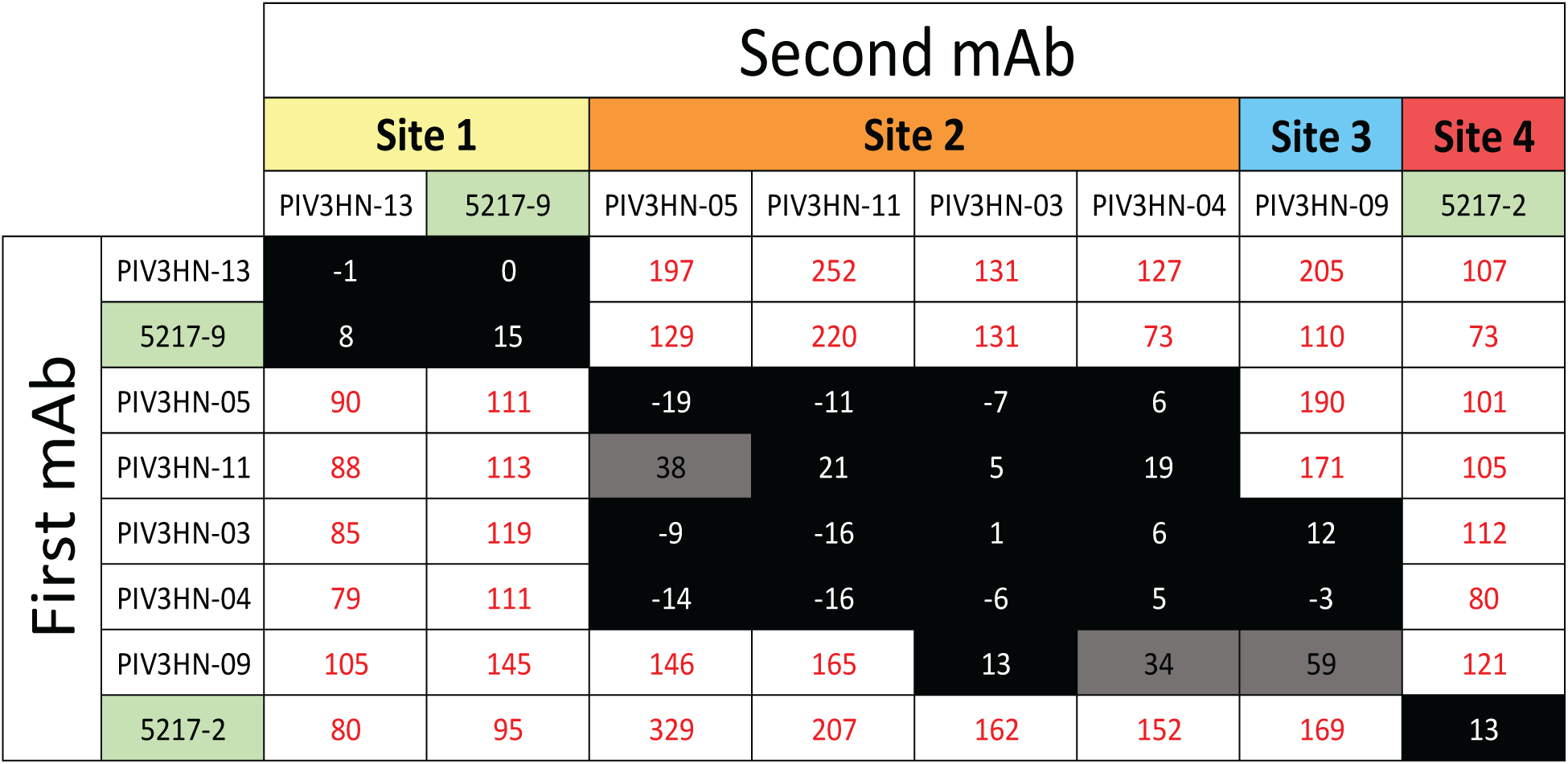
Biolayer interferometry (BLI) epitope binning analysis of novel mAbs against previously characterized mAbs. A heat map showing the results from a binning experiment with the six of the previously characterized mAbs (PIV3HN-13, PIV3HN-05. PIV3HN-11, PIV3HN-03, PIV3HN-04, PIV3HN-09) and the two new mAbs (5217-2 and 5217-9). Black indicates complete competition (≤33%), gray indicates partial competition (34-66%), and white indicates no competition (≥67%). The formula to calculate the competition percentage is as follows %Competition = (signal from first mAb with second mAb/signal from first mAb) x 100%. The addition of mAb 5217-2 led to the discovery of a new epitope on PIV3 HN, designated as Site 4, while mAb 5217-9 bound to the previously characterized Site 1. Both epitopes are highlighted in green.

#### Site 3 mAb PIV3HN-09 binds opposite of Site 1 mAb PIV3HN-13

While the structures of mAbs targeting Sites 1 and 2 have been previously elucidated, we sought to further define the neutralizing epitopes on the PIV3 HN protein, including antigenic Site 3 and the newly discovered Site 4. The Site 3-directed mAb, PIV3HN-09, was previously identified as a neutralizing mAb capable of inhibiting viral fusion and reducing PIV3 lung viral titers in a hamster infection model^46^. We determined the structure of mAb PIV3HN-09 in complex with the PIV3 HN protein using cryo-electron microscopy (cryo-EM). The Fab of PIV3HN-09 was complexed with the PIV3 HN protein (along with the previously discovered Site 1 mAb PIV3HN-13) and visualized using single particle cryo-EM, allowing us to resolve the binding interface at high resolution. The final electron density map was resolved at a resolution of 2.95 Å obtained from 198,000 particles, and a predicted model from AlphaFold3 was fit into the map **(Figure 4A, 4B)**. Manual model building was performed using Coot, in which amino acid side chains and backbone conformations were adjusted to fit the cryo-EM density map. After model adjustment, refinement and validation were carried out in Phenix, which included evaluation of stereochemistry and Ramachandran outliers **(Table S1, Figure S2)**. Structural analysis reveals that mAb PIV3HN-09 binds at the opposite end of the Site 1 mAb, PIV3HN-13, near the HN dimer interface **(Figure 4A-C)**. Closer examination of the binding interface shows that the epitope is primarily recognized through interactions mediated by the heavy chain of the antibody. All three complementarity-determining region (CDR) loops of the heavy chain contribute to binding by engaging multiple PIV3 HN surface residues. Specifically, the HCDR1 residue Tyr38 forms a hydrogen bond with His231 of the PIV3 HN protein at a distance of 2.8 Å. Ser59 at the HCDR2 interacts with Pro232 (3.6 Å) of the PIV3 HN protein via a hydrogen bond. Tyr106 on the HCDR3 forms separate hydrogen bonds with Ser171 and Ser172 (2.3 Å and 3.1 Å, respectively). A final HCDR3 residue Lys108 forms a bond with Glu230 on HN (3.4 Å) **(Figure 4C, 4D)**. Notably, the light chain of mAb PIV3HN-09 does not appear to contribute to antigen binding, as no significant contacts are observed between light chain residues and the HN protein.

**Figure 4.**
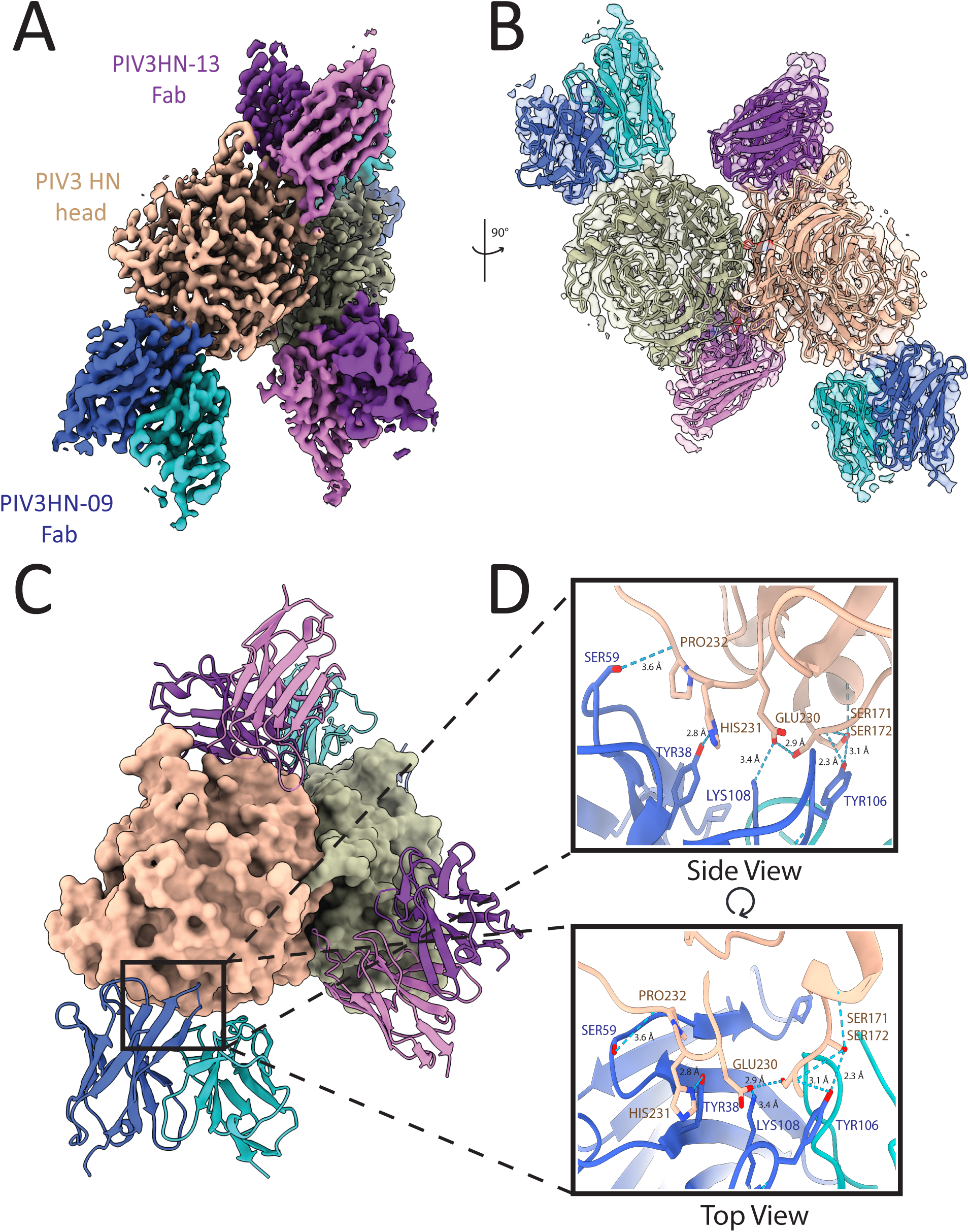
Cryo-EM structure of Site 3 mAb PIV3HN-09 binding to the PIV3 HN head. (A) Cryo-EM density map of a PIV3 HN dimer (tan & olive green) bound to PIV3HN-09 Fabs (blue) and PIV3HN-13 Fabs (purple). (B) Alternative view of the atomic model (ribbon representation) fitted into the cryo-EM density (transparent surface), demonstrating a good model to map fit. (C) Surface representation of the HN proteins with bound Fab fragments, PIV3HN-09 heavy chains are colored dark blue and light chains are light blue. (D) Detailed view of the interaction interface between the HN protein and the PIV3HN-09 heavy chain, highlighting key interactions. Hydrogen bonds are depicted as dotted lines, with distances labeled in angstroms (Å). The top panel shows a side view of the binding interface, while the bottom panel provides a top-down perspective.

#### Site 4 mAb 5217-2 binds opposite of the F-HN binding interface

To complete the structural characterization of the four known antigenic sites on the PIV3 HN surface, we utilized cryo-EM to resolve the Site 4 epitope interactions of mAb 5217-2. In this case, we leveraged the now known epitope of mAb PIV3HN-09 by complexing this mAb and mAb 5217-2 with the PIV3 HN protein. We obtained a 3.29 Å resolution electron density map from 238,000 particles **(Figure 5A, S3, Table S1)**. Structural analysis revealed that the mAb 5217-2 binds to the apex of the PIV3 HN head domain, positioned distal to the F-HN binding interface **(Figure 5A, 5B)**. Two heavy chain contacts predominantly drive the interaction, with an additional contribution from a single light chain residue. Unlike the PIV3HN-09 epitope, which engages multiple CDR loops, binding of the 5217-2 Fab relies largely on a single CDR loop. HCDR3 residue Asp108 forms two stabilizing hydrogen bonds with Arg62 (3.3 Å) and Lys60 (2.8 Å). Additional support for the interaction comes from framework residues: HFR3 residue Asp66 forms a hydrogen bond with Lys57 (3.3 Å), and LFR3 residue Asn66 forms a hydrogen bond with Leu 411 (3.5 Å) **(Figure 5D)**.

**Figure 5.**
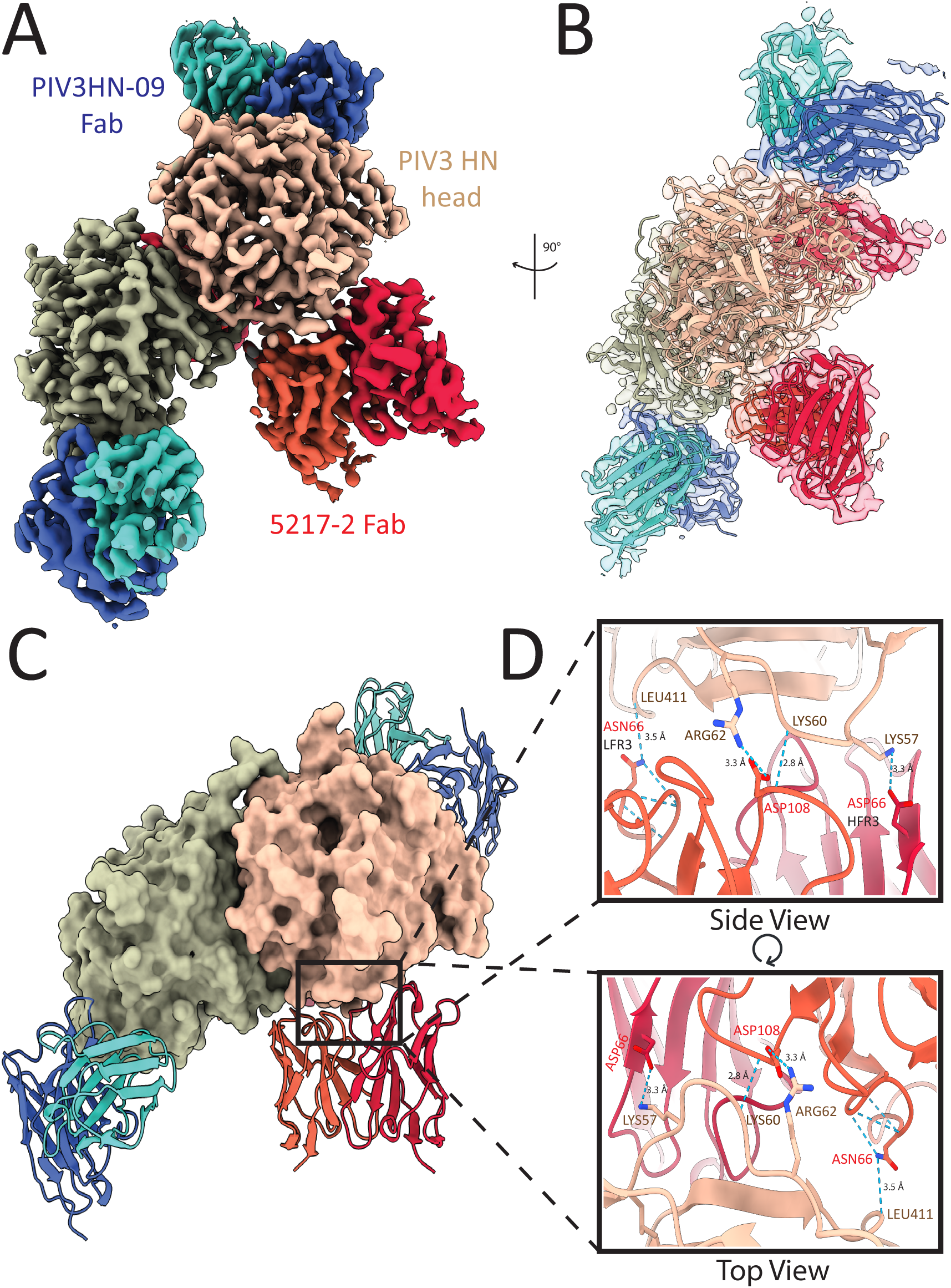
Cryo-EM structure of Site 4 mAb 5217-2 binding to the PIV3 HN head. (A) Cryo-EM density map of a PIV3 HN dimer (tan & olive green) bound to 5217-2 Fabs (red) and PIV3HN-09 Fabs (blue). (B) Alternative view of the atomic model (ribbon representation) fitted into the cryo-EM density (transparent surface), demonstrating a good model to map fit. (C) Surface representation of the HN proteins with bound Fab fragments, 5217-2 heavy chains are colored dark red and light chains are orange. (D) Detailed view of the interaction interface between the HN protein and the 5217-2 heavy and light chains, highlighting key contact residues. Hydrogen bonds are depicted as dotted lines, with distances labeled in angstroms (Å). The top panel shows a side view of the binding interface, while the bottom panel provides a top-down perspective.

### The molecular basis of cross-protection from PIV1 and PIV3 by the Site 2 mAb PIV3HN-05

The potent neutralizing activity of Site 2 mAbs is attributed to their targeting of the sialic acid binding site on the PIV3 HN protein, where they likely interfere with receptor binding and/or prevent viral release by blocking the interaction of the PIV3 HN protein with sialic acid^46^. Notably, the Site 2 mAb, PIV3HN-05, has demonstrated both prophylactic and therapeutic efficacy against PIV3 and exhibited cross-neutralization activity against PIV1, with an IC_50_ of 180 ng/mL^46^. To better understand the molecular basis for this cross-neutralization, we revisited the structural characterization of the mAb PIV3HN-05 epitope. The initial cryo-EM map had disordered density within the variable region, which limited the resolution and the elucidation of the key Fab-HN interactions. Using the updated CryoSPARC package, we were able to re-pick a larger number of particles and pool them with previously selected particles using improved templates. This enabled more effective 2D and 3D classification to remove resolution-limiting particles. It was also followed by a “rebalance orientation” job to lower the orientation dominancy of the preferred views. Applying the latest version of non-uniform refinement in CryoSPARC using a tighter manual mask ultimately allowed us to generate a higher-resolution cryo-EM map of the PIV3 HN protein in complex with PIV3HN-05 Fab and PIV3HN-13 Fab at 3.29 Å resolution from 343,000 particles, providing a clearer understanding of the mode of binding **(Figure 6A, S4, Table S1)**.

**Figure 6.**
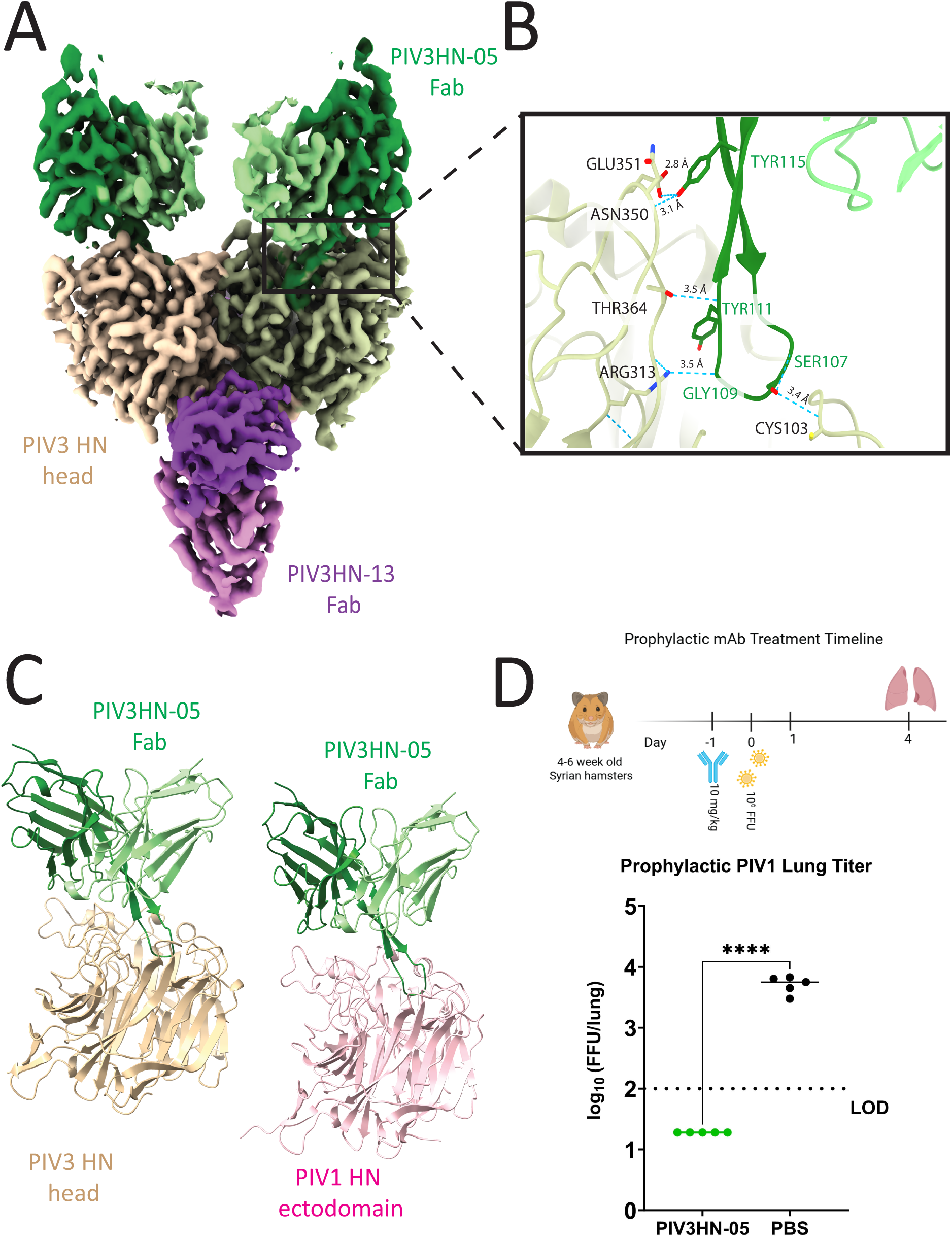
Updated cryo-EM structure of Site 2 mAb PIV3HN-05 binding to the PIV3 HN head and evaluation of cross-reactivity in a PIV1 viral challenge model. (A) Cryo-EM density map of a PIV3 HN dimer (tan & olive green) bound to PIV3HN-05 Fabs (green) and PIV3HN-13 Fabs (purple). (B) Detailed view of the interaction interface between the HN protein and the HCDR3 loop of PIV3HN-05 Fab. The Fab does not directly interact with the HN sialic acid active site (residues 234-323) but inserts into the active site, occupying the space. Hydrogen bonds are depicted as dotted lines, with distances labeled in angstroms (Å). (C) The left panel shows the overall structure of the PIV3 HN head domain in complex with the PIV3HN-05 Fab. To evaluate the potential mechanism behind PIV1 cross-reactivity, the PIV1 HN ectodomain was structurally aligned with the PIV3 HN head, and the binding site of the PIV3HN-05 Fab was compared between the two. (D) In vivo evaluation of PIV3HN-05 against PIV1 challenge. Top: Experimental timeline of the hamster study. Syrian hamsters (n=10) received a single dose of PIV3HN-05 (10 mg/kg) one day prior to the PIV1 viral challenge. Lungs were harvested four days post-challenge, and viral titers were assessed via plaque assay. Bottom: Results of the plaque assay show a reduction in lung viral load in the PIV3HN-05 treated group compared to the PBS control. Statistical significance was determined using a two-tailed unpaired t-test, revealing a highly significant difference between groups (P < 0.0001). Animals with undetectable viral titers were plotted as the limit of detection (LOD) divided by 2.

The structure revealed that the interaction between PIV3HN-05 Fab and the PIV3 HN protein is mediated exclusively by the heavy-chain complementarity-determining region 3 (HCDR3) loop. This loop fits deeply within the binding pocket of the PIV3 HN sialic acid binding site, likely limiting the ability of the protein to bind to the host cell sialic acid-containing receptors. This interaction is mediated primarily by five hydrogen bonds between the HCDR3 of the Fab and HN active site. Residue Tyr115 forms two stabilizing hydrogen bonds with Glu351 (2.8 Å) and Asn350 (3.1 Å), Tyr111 interacts with Thr364 (3.5 Å), Ser107 interacts with Cys103 (3.4 Å), and Gly109 interacts with Arg313 (3.5 Å) **(Figure 6B)**. These contacts suggest a mechanism in which the Fab physically blocks the sialic acid binding site, therefore inhibiting viral attachment and entry into the host cell. Since we previously showed PIV3HN-05 cross-neutralized PIV1, we overlaid the HCDR3 loop onto PIV1 HN to assess the mechanism behind the cross-reactivity for PIV1 **(Figure 6C)**^46^. The HCDR3 loop appears to insert into the sialic acid binding pocket of PIV1 HN, suggesting a similar mechanism of neutralization against both PIV1 and PIV3.

To further evaluate the protective efficacy of the PIV3HN-05 mAb against PIV1, we conducted an in vivo prophylactic challenge study. For this study, we utilized Syrian golden hamsters as the animal model, as they support PIV1 replication in both the lung and nasopharynx. Hamsters received an intraperitoneal injection of mAb PIV3HN-05 at a dose of 10 mg/kg or PBS as a control, administered 24 hours before the intranasal PIV1 challenge. Four days after the challenge, animal lung were harvested and homogenized for viral titration via plaque assay **(Figure 6D, top)**. Compared to the PBS-treated group, which exhibited an average of 6.8 x 10^3^ ffu/mL per lung, the PIV3HN-05 mAb-treated group showed complete inhibition of viral replication, with no detectable plaques at the limit of detection, indicating effective reduction of PIV1 in the lungs **(Figure 6D, bottom).** These findings indicate that mAb PIV3HN-05 provides protection not only against PIV3 but also against PIV1, likely due to its ability to block HN binding to host cell receptors through a conserved mechanism of neutralization.

#### Site 1-4 mAbs bound to the PIV3 HN head

To gain further insight into the potential mechanism of neutralization, we fit the structures of the PIV3 HN-specific mAbs onto the PIV3 F-HN complex cryo-ET map (EMD-27550), which captured the interactions of the PIV3 F and HN proteins on authentic viral particles **(Figure 7A, 7B)**^52^. This structural alignment revealed that mAb PIV3HN-09 binds adjacent to the F-HN interaction site. The location of this binding site is consistent with previous functional data showing that mAb PIV3HN-09 inhibits fusion between the PIV3 F and HN proteins^46^. Comparison of the Site 4 binding position relative to the F-HN interface reveals that it is situated adjacent to Site 2 **(Figure 7A, 7B)**. Site 2-specific mAbs bind within the sialic acid-binding pocket of the PIV3 HN protein, whereas structural alignment demonstrated that the Site 4 mAb binds just outside this pocket, adjacent to the receptor-binding region. Despite not directly overlapping the sialic acid binding site, the Site 4 antibody 5217-2 exhibited potent neutralizing activity.

**Figure 7.**
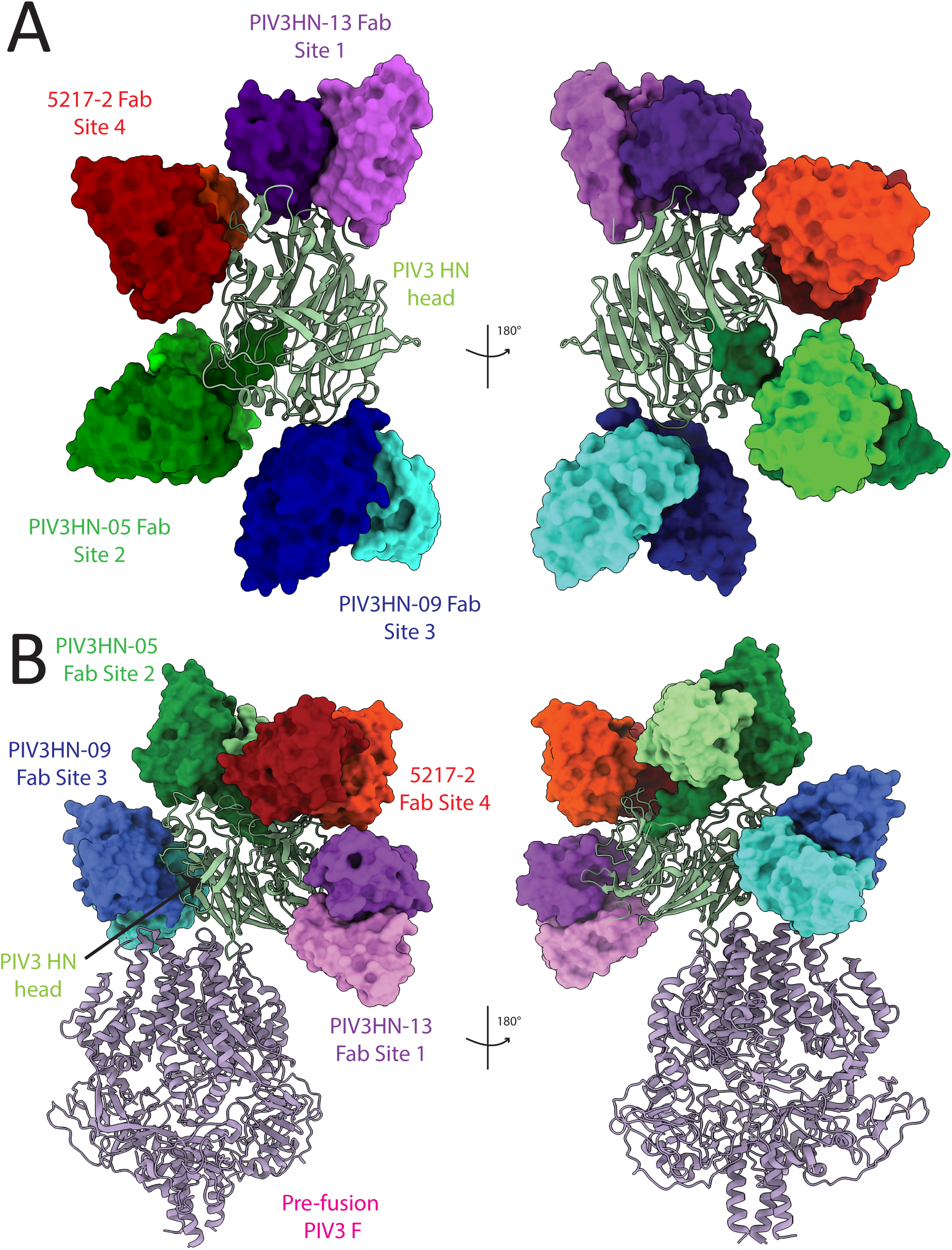
Structural mapping of Site 1-4 Fabs bound to the PIV3 HN head domain in complex with the PIV3 F protein. (A) A model showing the PIV3 HN head domain simultaneously bound by the four site-specific Fabs: Site 1 PIV3HN-13, Site 2 PIV3HN-05, Site 3 PIV3HN-09, and Site 4 5217-2. This model provides a comprehensive representation of the epitope location on the HN head, illustrating the relative position and accessibility of each site. (B) A model of the PIV3 HN head domain with Site 1-4 Fabs bound in the context of the PIV3 F protein, generated using a structure of prefusion F (PDB: 6MJZ) and the HN-F interaction map (EMB: 27550). This model shows the orientation and relative positioning of each Fab when the HN is engaged with the F protein.

#### Site 4 mAb 5217-2 prevents viral replication in Syrian hamster lungs

Given the proximity of mAb 5217-2 to the receptor-binding region on the PIV3 HN protein, together with its potent in vitro neutralization and complement deposition activity, we next evaluated whether this mAb could confer protection in vivo in a prophylactic setting. Syrian golden hamsters were treated 24 hr before infection with 10 mg/kg of mAb or PBS control via intraperitoneal injection. Hamsters were then challenged with PIV3 intranasally, lung samples were collected, and viral load in the lungs were quantified via plaque assay **(Figure 8A)**. Administration of mAb 5217-2 resulted in a significant reduction in viral burden compared to the control-treated animals. PBS-treated hamsters exhibited a mean viral load of 2.2 x 10^4^ ffu/lung, whereas animals treated with mAb 5217-2 showed significantly lower titers, averaging 4.7 x 10^3^ ffu/lung **(Figure 8B)**. This represents approximately a 4.5-fold reduction in viral replication within the lung.

**Figure 8.**
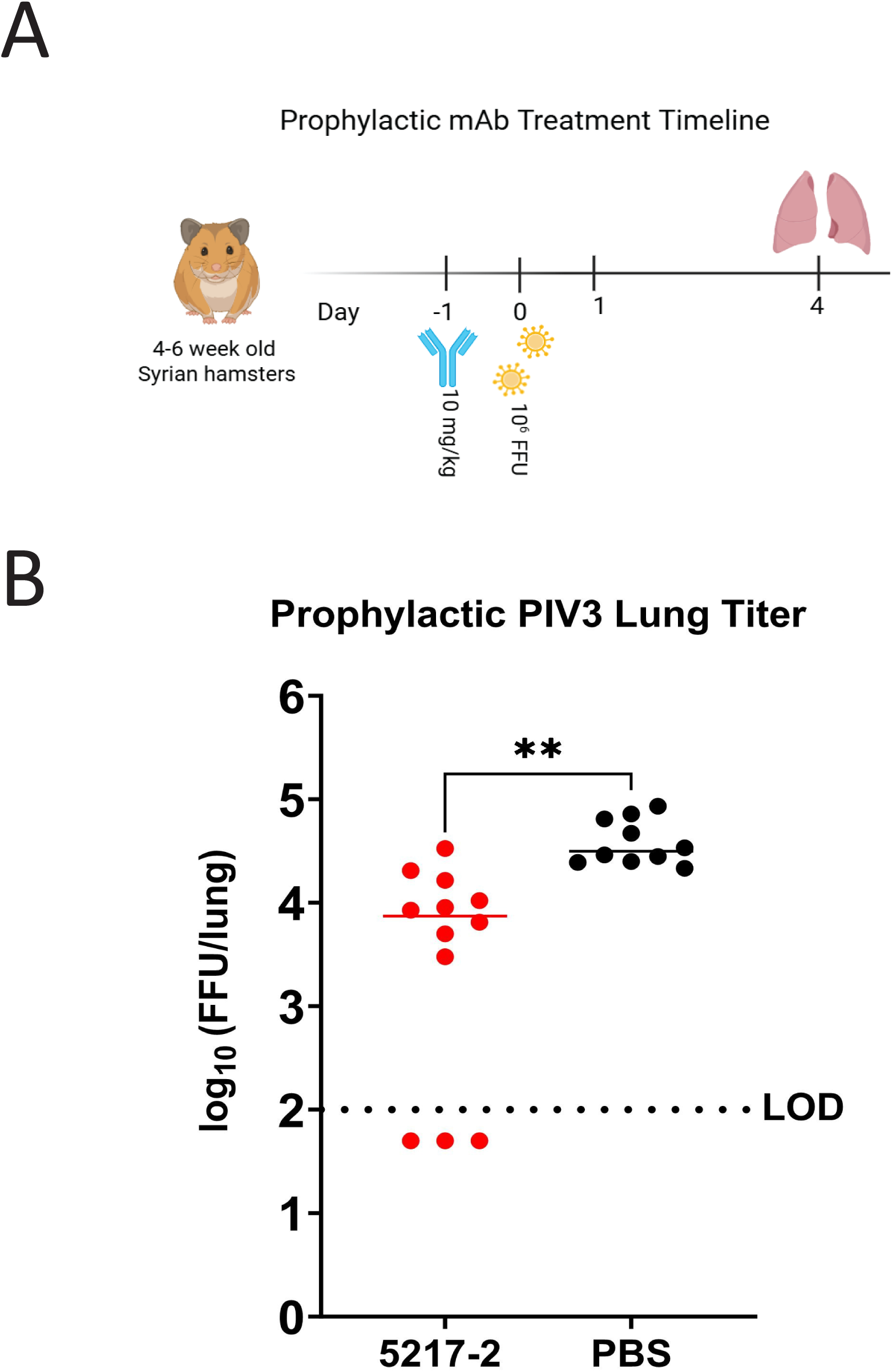
In vivo evaluation of 5217-2 against PIV3 challenge. (A) Experimental timeline of the hamster study. Syrian golden hamsters were treated prophylactically with 5217-2 (10 mg/kg) or PBS 24 hr prior to intranasal PIV3 challenge. Lungs were harvested four days post-challenge, and viral titers were assessed via plaque assay. (B) Results represent pooled data from two independent experiments. Two animals were excluded from the analysis due to failed viral infection evidenced a plaque count of less than 10 in the 1/10 dilution well. Plaque assay analysis demonstrated a 1-log reduction in lung viral titers in the 5217-2 treated group compared to PBS controls. Statistical significance was determined using a two-tailed t-test, revealing a significant difference (P = 0.0032) between groups. Animals with undetectable viral titers were plotted as the limit of detection (LOD) divided by 2.

## Discussion

With no approved vaccines or antiviral therapies currently available for PIVs, there is a critical need for the development of effective preventative and therapeutic strategies. Neutralizing mAbs have shown clinical success against several respiratory viruses demonstrating their potential to mitigate disease severity in vulnerable populations. However, to date, no mAbs have been approved for the treatment or prevention of PIV infections, highlighting a significant gap in available interventions against this virus. In this study, we introduced two mAbs that target the PIV3 HN antigen and characterized the binding avidity, viral neutralization ability, in vivo prophylactic activity, and structural properties of these mAbs. With this, we were able to determine additional epitopes on the PIV3 HN protein and demonstrated its potential as a viable target for vaccines or mAb development.

With the addition of the fourth epitope, Site 2 of PIV3 HN remains the primary target of the most potent neutralizing mAbs. Although the Site 2 mAb PIV3HN-03 was previously reported as the strongest inducer of complement deposition among the panel, we observed that the Site 4 mAb 5217-2 exhibited increased complement deposition relative to mAb PIV3HN-03.^46^ In addition to the efficacy studies, we determined the structures of mAbs targeting Site 3 and Site 4 on the PIV3 HN protein to gain insight into the mechanism of protection. Structural analysis revealed that the Site 3-specific mAb, PIV3HN-09, binds adjacent to the interface between the PIV3 HN and F proteins. Given that viral entry depends on coordinated HN-F interactions to trigger F protein-mediated membrane fusion^22^, the position of PIV3HN-09 suggests that it interferes with this activation step. Consistent with its ability to inhibit viral fusion^46^, the orientation of PIV3HN-09 at the HN-F interface suggests that its binding imposes steric constraints that limit efficient HN-mediated activation of the F protein.

Structural reanalysis of the PIV3 HN protein-mAb PIV3HN-05 interaction reveals that the HCDR3 loop directly occupies the sialic acid binding pocket of the HN protein, potentially blocking the HN’s ability to bind to sialic acid on the host cell. Consistent with this binding mode, mAb PIV3HN-05 exhibits potent neutralizing activity and cross-reactivity with PIV1, underscoring the sialic acid binding pocket as a key functional target for HN-directed neutralization. The cross-reactive properties of mAb PIV3HN-05 are further supported by the high degree of structural conservation between the HN proteins of PIV1 and PIV3. In contrast, the structure of mAb 5217-2 revealed that the Site 4 epitope is positioned at the apex of the HN head domain, adjacent to the receptor-binding region targeted by mAb PIV3HN-05. Although the Site 4 epitope does not directly overlap this active site, its proximity to this functional region suggests that mAb binding at Site 4 could influence HN function through sterically limiting receptor access or by restricting the conformational flexibility of the HN head domain required for receptor binding. This proposed mechanism is supported by the observed reduction in viral load in vivo following prophylactic administration of mAb 5217-2. Together, these data suggest that epitopes adjacent to the receptor-binding site can mediate protective immunity, highlighting Site 4 as a functionally relevant region of PIV3 HN.

Several limitations of this study should be considered. First, the in vivo efficacy was evaluated exclusively in a prophylactic setting, prior to the establishment of PIV infection. As a result, it remains unclear whether these mAbs would be effective in a therapeutic context, which is an important factor in clinical applications. Second, although murine models are often an effective method of measuring antibody-mediated protection, species-specific differences in complement regulation between rodents and humans may influence Fc effector functions and limit the translational interpretation of these mAbs.^53^ Finally, Fc effector function was primarily tested through complement-induced cytotoxicity; however, mAbs can also engage additional Fc-dependent mechanisms, such as recruiting innate immune cells to induce phagocytosis or NK-mediated cytotoxicity. Together, these limitations highlight the need for future studies incorporating therapeutic challenge models and a more comprehensive evaluation of Fc-mediated effector mechanisms to further define the protective efficacy of HN-targeting antibodies.

In summary, our study offers comprehensive new insights into the antigenic landscape of the PIV3 HN protein, highlighting the distinct roles of various epitopes in mediating antibody protection. The identification of multiple antigenic sites on PIV3 HN with distinct functional outcomes provides a framework for rational vaccine design strategies that elicit antibodies targeting conserved and critical regions, such as Site 2. By integrating epitope binning, plaque reduction neutralization assays, structural characterizations, and in vivo efficacy studies, we concluded that mAbs targeting Site 2 and Site 4 are the most promising targets for further investigation. Our findings emphasize the robust protective capacity of Site 2-targeting antibodies, particularly mAb PIV3HN-05 which binds a critical and conserved region on the PIV3 HN protein. To further understand the basis for its cross-protective activity, future studies should investigate the structural interactions between mAb PIV3HN-05 and the HN protein of PIV1. Additionally, further studies should be done to evaluate the in vivo efficacy and structural interactions of other Site 2 mAbs, such as PIV3HN-11, which was shown to inhibit fusion of PIV3 to host cells^46^. Together, these efforts will help guide the development of broadly protective monoclonal antibodies targeting the HN protein of PIV3, as well as those from other parainfluenza viruses.

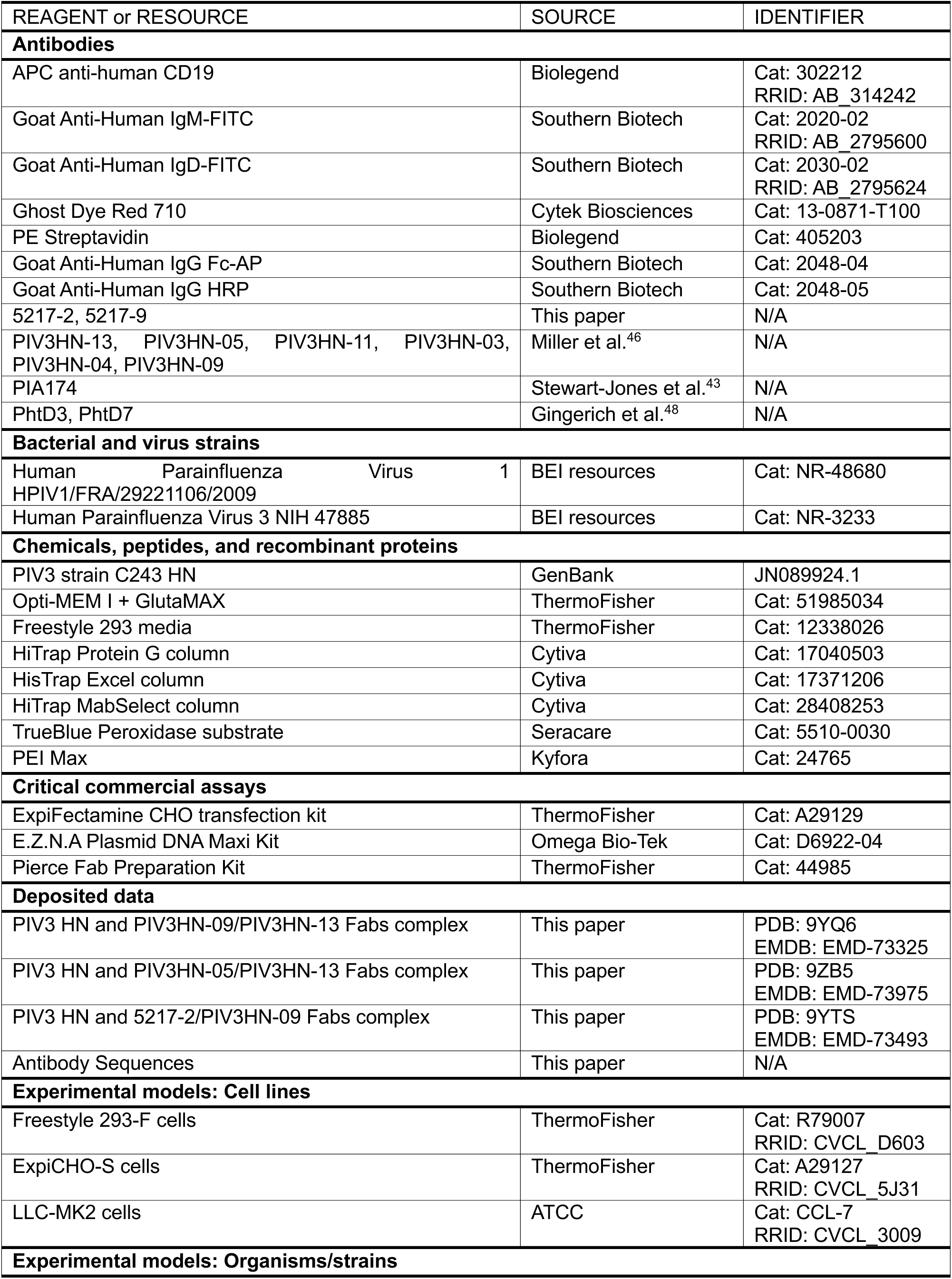
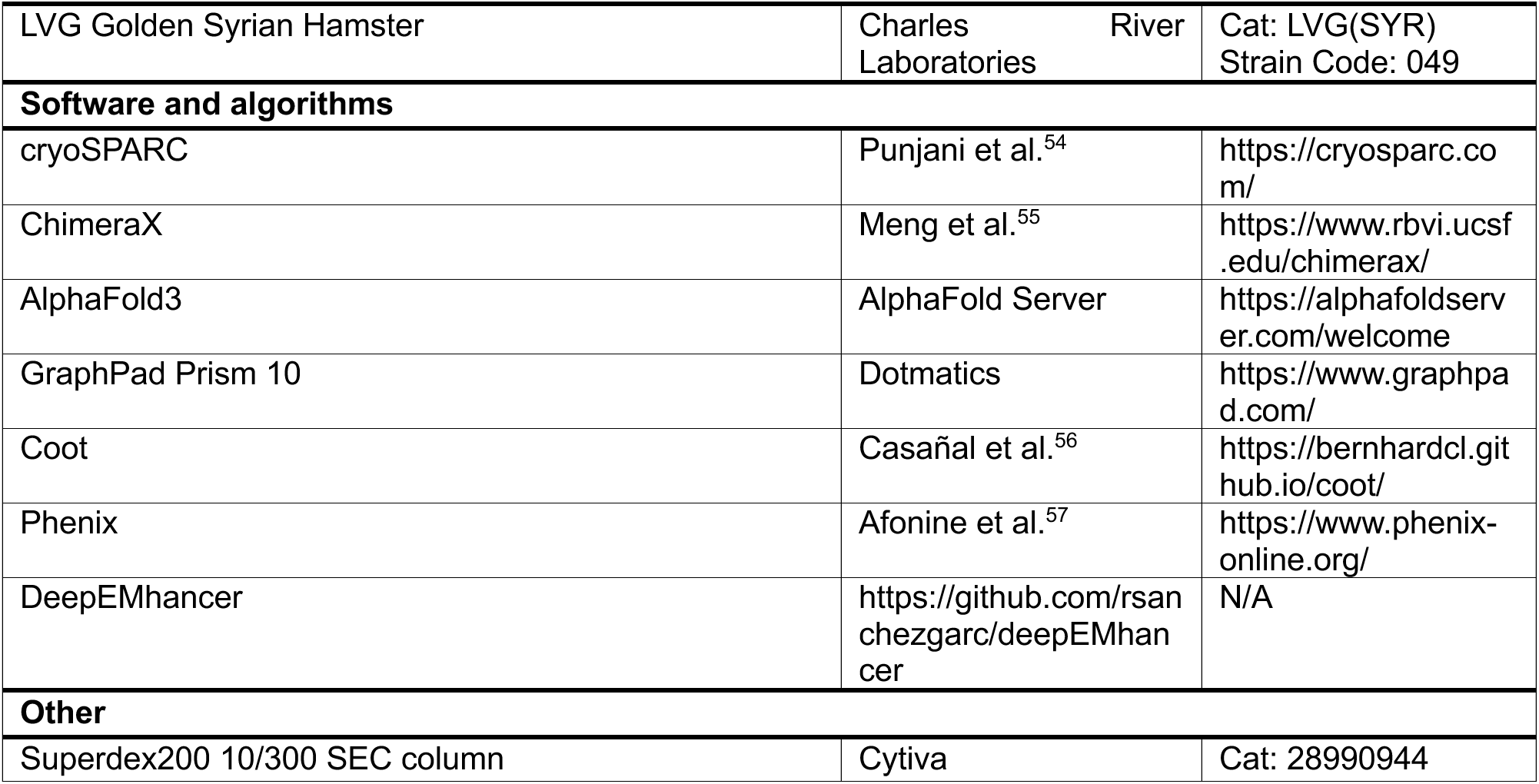

## Methods

### Cell lines and viruses

Freestyle293-F cells (ThermoFisher, Cat: R79007) were cultured in Freestyle 293 media (ThermoFisher, Cat: 12338026). ExpiCHO-S cells (ThermoFisher, Cat: A29127) were cultured in ExpiCHO Expression Medium (ThermoFisher, Cat: A2910001). LLC-MK2 cells (ATCC, CCL-7) were cultured in Opti-MEM I + GlutaMAX (ThermoFisher, Cat: 51985034) supplemented with 2% fetal bovine serum. PIV1 (Cat: NR-48680) and PIV3 (Cat: NR-3233) was obtained from BEI resources. The virus was diluted into virus medium (Opti-MEM, 1:500 trypsin-EDTA, 1:100 CaCl_2_, and 1% antibiotic-antimycotic) and added to LLC-MK2 cells at an MOI of 0.01. Virus was propagated at 5% CO_2_ for 3–5 days until CPE was observed. The virus was subsequently harvested using a sucrose freeze/thaw method. The virus-containing sucrose solution was aliquoted, flash-frozen in liquid nitrogen, and stored at − 80 °C until further use.

#### Production of PIV3 HN viral protein

A plasmid construct for the head domain of PIV3 HN was designed based on the wild-type PIV3 strain C243 HN sequence (GenBank JN089924.1) with the N-terminal cytoplasmic tail and transmembrane domains removed and a Hexahistidine tag added. Plasmid containing the HN head domain insert was transformed into DH5a E. coli competent cells prior to plasmid isolation and transfection. The recombinant protein was expressed in Freestyle293-F cells. Plasmids were diluted in Opti-MEM medium at a final concentration of 1 µg/ml. The transfection reagent PEI MAX (Kyfora Bio, Cat: 24765) was added at a ratio based on the culture volume and incubated with the plasmid mixture for 30 mins before adding to the cells. Cultures were centrifuged at 10,000 RPM for 15 mins and then the supernatant was passed through a 0.45 µm filter. Protein was purified through a HisTrap Excel pre-packed column (Cytiva, Cat: 17371206) per the manufacturer’s instructions. Purified protein was buffer exchanged into PBS and stored at − 80 °C until use.

#### Peripheral blood mononuclear cells (PBMCs) Isolation

Leukocyte filters obtained from Shepeard Blood Bank (Augusta, GA) were emptied into sterile tubes and flushed twice with phosphate-buffered saline (PBS). The collected blood was carefully layered on top of pre-warmed Lymphocyte Separation Medium (Corning, Cat: 25-072-CV) and then centrifuged at 2500 RPM for 30 mins using minimal brake and acceleration (set at 1) to not disturb the layers. The plasma was removed, and the buffy coat was carefully extracted and added to Dulbecco’s Modified Eagle Medium (DMEM) (Corning, Cat: 10-013-CV). The PBMCs were washed twice with DMEM (300 x g for 15 mins) and PBMCs were aliquoted into cryovials with Stemcell Media A (Stemcell Technologies, Cat: 03801) and 10% DMSO (ThermoScientific, Cat: J66650.AE) was added to minimize cell damage during freezing. Cryovials were placed in a cryogenic freezing container and stored at –80°C overnight. For long time preservation, samples were transferred to liquid nitrogen storage.

### 5217-2 and 5217-9 mAb Discovery

PBMCs from blood of human donors were blocked with Trustain FcX (Biolegend, Cat: 422302) for 30 minutes, washed once in PBS, and subsequently stained according to the following protocol: 5 µL CD19-APC (Biolegend, Cat: 302212), 1.25 µL IgM-FITC (Southern Biotech, Cat: 2020-02), 1.25 µL IgD-FITC (Southern Biotech, Cat: 2030-02), 2.5 µL GhostRed710 (Cytek Biosciences, Cat: 13-0871-T100), and 5 µL of PIV3 HN conjugated to PE (Biolegend, Cat: 405204). The cells were stained for 30 minutes, washed twice in PBS, and immediately sorted on Moflo Astrios EQ (Beckman Coulter). Prior to antigen-specific sorting, 9 pneumococcal proteins (PcpA, PspA, PsaA PhtE, Ply, NanA, PcsB, StkP, PiuA) and 2 viral proteins, PIV HN and RSV B2F Pre, were biotinylated using the EZ-Link NHS-PEG4 Biotinylation kit as per the manufacturer’s protocol (ThermoFisher, Cat: 21455). Streptavidin-conjugated PE (Invitrogen, Cat #S866) was slowly added to the biotinylated antigens at a fluorophore to protein molar ratio of 4:1, adding 1 uL of streptavidin-PE every 20 minutes for a total of 5 additions and stored on ice away from light. PBMCs were suspended in FACS buffer (PBS, 2% FBS) (Gibco, Cat # A5256701), 2% goat serum (Gibco, Cat #16210072), 0.5M EDTA) and Fc-blocked with Human TruStain FcX (BioLegend, Cat #422301) on ice for 30 min. The sorted cells were processed through 10X sequencing (Pleasanton, CA) at the UGA Genomics and Bioinformatics Core, according to the manufacturer’s protocol (10XGenomics, Protocol: CG000736). After 10X processing and analysis, paired heavy and light chain variable sequences were obtained **(Table S2)** and cloned by Twist Biosciences (San Fransicsco, CA) into their respective cloning vector (heavy: pTwist CMV hIgG1, lambda: pTwist CMV hIgL2, kappa: pTwist CMV hIgK).

#### Plasmid transformation into E. coli competent cells

Plasmids from 10x sequences were ordered from Twist Biosciences and transformed into *Escherichia Coli* DH5α competent cells (Invitrogen, Cat: 18258012). The cells were incubated with plasmid DNA (1 µL) and heat shocked before incubating in LB broth medium for 1 hr and plated on ampicillin agar plates overnight. Isolated colonies were then added to a 5 mL culture of LB broth with ampicillin (1:1000) and then expanded to a 200 mL culture 8 hours later to shake overnight. The DNA plasmids were extracted using the E.Z.N.A Plasmid DNA Maxi Kit (Omega Bio-Tek, Cat: D6922-04) according to the manufacturer’s instructions. The resulting plasmid DNA was sterile filtered for later use in transfections.

#### Plasmid transfection into Freestyle 293-F cells and ExpiCHO cells

Both Freestyle 293F cells and ExpiCHO-S cells were used to transfect the PIV mAbs. In vitro studies utilized mAb produced from ExpiHO-S cells, while the in vivo studies used mAb produced from Freestyle293F cells. For Freestyle 293F transfections, heavy and light chain plasmids were diluted in Opti-MEM medium at a final concentration of 1 µg/ml. The transfection reagent PEI MAX (Kyfora Bio, Cat: 24765) was added at a ratio based on the culture volume and incubated with the plasmid mixture for 30 mins before adding to the cells. Transfections using ExpiCHO-S cells (ThermoFisher, Cat: A29127) were performed according to the ExpiFectamine CHO transfection kit (ThermoFisher, Cat: A29129) based on the Max Titer Protocol.

#### Purification of mAbs

mAbs were purified seven days post-transfection in Freestyle 293F cells or fourteen days post-transfection in ExpiCHO-S cells. Cultures were centrifuged at 10,000 RPM for 15 mins and then the supernatant was passed through a 0.45 µm filter. A HiTrap Protein G column (Cytiva, Cat: 17040503) was used to purify the supernatant according to the manufacturer’s instructions. The eluted mAbs were spun in 30 kDa MWCO concentrators (Millipore, Cat: UFC903096) and buffered exchanged with PBS. The concentration was measured on a NanoDrop spectrophotometer using the IgG setting.

#### Enzyme-linked Immunosorbent Assay (ELISA)

To measure binding to the PIV3 HN protein, 384-well plates were coated with 2 µg/ml of antigen overnight at 4 °C. The plates were blocked for 1 hr with 2% non-fat dry milk supplemented with 2% goat serum. Plates were then washed three times with water and primary mAbs were applied to wells for 1 hr at 37 °C. Following another three times wash with water, a secondary antibody goat anti-human IgG Fc-AP (Southern BioTech, Cat: 2048-04) at a dilution of 1:4000 in 1% blocking solution was added. After incubating for an hour, the plates were washed five times with PBS-T (0.05% Tween-20 in PBS) and substrate solution (1 mg/ml PNPP disodium salt hexahydrate, ThermoFisher) was added to each well. The plates were incubated in the dark for 1 hr at room temp before reading the optical density at 405 nm on a Biotek plate reader.

#### Complement deposition cytotoxicity (CDC) assay

Biotinylated PIV3 HN antigen was coupled with FluoSpheres Neutravidin beads (Invitrogen, Cat: F8775) at a 1:1 ratio of antigen (µg): beads (µL) for 2 hr at 37°C. The beads were washed with 5% PBS-BSA and resuspended 1:100 in 0.1% PBS-BSA. The mAbs were diluted to 2.5 µg/mL in a 96 U-bottom plate and incubated for 2 hr with 10 µL of antigen-specific beads. The bead-mAb complexes were then washed with 0.1% PBS-BSA. Guinea pig complement (MP Biomedicals, Cat: 64283) was diluted 1:50 in R-10 buffer (RPMI-1640+10% FBS). The diluted complement was then incubated with the bead-mAb complex for 15 mins and then washed twice with PBS. Fluorescein-conjugated goat anti-guinea pig complement c3 (MP Biomedicals, Cat: 55385) was diluted 1:100 in PBS and incubated for 15 min in the dark. Complexes were washed twice with PBS and resuspended in a final volume of 150 µL PBS. Samples were read on the Cytek Aurora.

#### Flow cytometry with PIV3-infected cells

Biotinylated mAbs were incubated with streptavidin-conjugated PE (BioLegend, Cat: 405204) on ice for 1 hr and stored at 4°C for later use. LLC-MK2 cells were cultured in a T225 flask at 37°C, 5% CO_2_ to 80-90% confluency prior to infection. Cells were washed with PBS twice and infected with PIV3 at an MOI of 0.01. The virus was incubated with the cells for 1 hr at 37°C, 5% CO_2_ with gentle rocking every 10 mins. Virus culture media (OptiMEM + 5 µg/mL trypsin-EDTA + 100 µg/mL CaCl_2_ + 1% 100x anti-anti) was added to the flask and incubated for 48 hrs. After incubation, the cells were digested using Versene (Gibco, Cat:15040-066) at 37°C, 5% CO_2_ for 45 mins. Cells were washed twice with FACS buffer and incubated in FACS for 30 mins on ice. Aliquots of 1 million cells were incubated with their respective PE-conjugated mAbs at 2.5 µg/mL for 30 mins on ice in the dark. Cells were again washed twice with FACS buffer and fixed for 15 mins with 4% paraformaldehyde. Finally, the cells were resuspended in 1 mL of FACS buffer. Samples were read on the Cytek Aurora.

#### PIV3 mAb plaque reduction neutralization assay

LLC-MK2 cells were seeded in 24-well flat bottom plates and incubated at 37°C, 5% CO2 for 2 days to reach confluency. Following the incubation period, 5217-2 and 5217-9 mAbs were serially diluted and incubated at a 1:1 ratio with PIV3 for 1 hr at room temperature. The media was removed from the 24-well plates and 50 µL/well of the virus/mAb mixture was added on top of the cells and rocked at 37°C, 5% CO2 for 1 hr. Cells were then overlaid with of 0.75% methylcellulose dissolved in Opti-MEM with 5 µg/mL trypsin-EDTA and 100 µg/mL CaCl_2_ and incubated for 4 days. After incubation, the cells were fixed with 10% neural buffered formalin for 45 mins and washed once with water. The plaques were immuno-stained by first blocking the plates for 1 hr with block comprised of 2% non-fat milk and 2% goat serum in PBS-T. The plates were washed 2 times with water and the primary mAb, PIA174, was then added at a dilution of 1 µg/mL in block and incubated for 1 hr at room temp. After washing 2 times with water, HRP-conjugated goat anti-human IgG (Southern BioTech, Cat: 2048-05) diluted 1:2000 in block was added for an hour. The plates were washed 3 times with water and TrueBlue peroxidase substrate (Seracare, Cat: 5510-0030) was added to each well and rocked for no longer than 10 mins and washed once with water. The plates were allowed to dry overnight and were manually counted using a stereoscope. IC_50_ values were calculated using GraphPad Prism10.

#### Epitope Binning

Biolayer interferometry on a Gator Prime instrument was performed for epitope binning. Anti-His (HIS) Probes (GatorBio, PN:160009) were equilibrated in kinetics buffer. A baseline measurement was recorded by dipping the biosensors into wells containing the kinetics buffer for 60 s. Next, biosensors were loaded with PIV3 HN at a concentration of 100 µg/mL for 150 s followed by another baseline measurement. The loaded biosensors were then dipped into wells containing either mAb PIV3HN-01, PIV3HN-03, PIV3HN-04, PIV3HN-05, PIV3HN-09, PIV3HN-11, PIV3HN-13, 5217-2, or 5217-9 at a concentration of 100 ug/ml for 300 s. The biosensors were then dipped into another set of the same mAbs listed previously for 300 s to test for a second association. Each antibody was tested for competition against itself and all other antibodies, with assays performed in both directions by using each antibody in both the first and second association steps.

#### Expression and purification of Fabs

Fab fragments PIV3HN-01, PIV3HN-09, PIV3HN-13, and 5217-2 were generated by incubating the full length mAbs with immobilized papain overnight at 37 °C, following the Pierce Fab Preparation Kit (ThermoFisher, Cat: 44985). The Fc portion was removed using HiTrap MabSelect columns (Cytiva, Cat: 28408253) according to the manufacturer’s instructions.

### Production of antigen + Fab complexes and Cryo-EM grid preparation

Fab fragments (0.7 mg/mL) were combined with PIV3HN (0.35 mg/mL) at a twofold molar excess and incubated overnight at 4 °C. The complex was purified on a Superdex200 10/300 SEC column (Cytiva, Cat: 28990944) with 50 mM Tris-HCl pH 7.5 and 150 mM NaCl. The purified complex peak was confirmed using SDS-PAGE gel stained with Coomassie blue and concentrated to 1 mg/mL for further use. A FEI Vitrobot MarkIV was used to prep sample onto Quantifoil R 1.2/1.3 300 copper mesh grids (Ted Pella, Cat: 658-300-CU). The grids were glow-discharged before adding 5 μL of sample and fast incubated before being blotted and plunged into liquid ethane.

#### Cryo-EM data collection & processing

After screening the prepared grids, 4000 movies (PIV09 complex) and 6300 movies (5217-2 complex) were collected on Titan-Krios 300KV microscope equipped with Gatan-K3 camera (Sup **Table S1** for details). Movies then imported into CryoSPARC package^54^ where patch motion correction and patch CTF estimation were performed. Blob picker was used to pick particles, which then were purified using multiple 2D classification cycles. Selected particles then were used to make multiple ab-initio 3D reconstruction which were used for further 3D classification (heterogenous refinement in CryoSPARC) to obtain the final particle set. Final 3D reconstruction was performed using the non-uniform refinement job to obtain a 2.95 Å (PIV09 complex) and a 3.29 Å (5217-2 complex) resolution map. We used the sharpened map from this refinement for model building and model refinement, and the sharpened map from DeepEMhancer for visualizations and figures.

#### Model building and validation

PIV3 HN dimer model and Fab aa sequences were all imported into Alphafold3^58^. After fitting and fixing the backbone and sidechains with precision into the map using Coot^56^; PHENIX^57^ was used to do a 3D real-space refinement and validation **(Table S1)**. Output models were used to fit into the CryoEM map using ChimeraX^55^.

#### Prophylactic/therapeutic treatment and viral challenge of Syrian hamsters

Four- to six-week-old female Golden Syrian hamsters (Charles River Laboratories #049) were individually housed at Florida State Universities’ animal facility. One day before the viral challenge, groups were prophylactically treated with their respective mAb (10 mg/kg) through intraperitoneal (IP) injection. After 24 hr, hamsters were anesthetized with isoflurane for 2 min before intranasally (IN) infecting with PIV3 (10^6^ PFU/mL). Hamsters were humanely euthanized using pentobarbital injection 4-days post-infection. Lungs were collected and washed with PBS and added to 2 mL of cold Opti-MEM and homogenized for viral titration. If not used immediately, lung samples were stored at −80°C before homogenization.

#### Syrian hamster lungs viral titration

After homogenization, the lung homogenate was serially diluted in cold Opti-MEM media. The diluted homogenate was added to LLC-MK2 cell monolayer in a 24-well tissue culture plate (200 µL/well) and rocked for 1 hr at 37°C, 5% CO_2_. The following steps of the titration assay were performed as described earlier.

## Acknowledgements.

This work was supported by the National Institute of Allergy and Infectious Diseases (NIAID) of the National Institutes of Health under award number R01AI143865 and R56AI181850 (to J.J.M.). The funders had no role in study design, data collection and analysis, decision to publish, or preparation of the manuscript. Florida State University supports cryo-EM in the Biological Imaging Resource Center, which houses the following equipment used in this study: a Gatan Solaris Plasma Cleaner (NIH grant S10 RR024564), a Hitachi HT7800 (NSF grant MRI2017869), a ThermoFisher Vitrobot Mark IV (NIH grant S10 RR024564), an SPI chameleon plunging system (NIH grant R24 GM145964), a ThermoFisher Titan Krios (NIH grant S10 RR025080), and a DE Apollo direct electron detector (NIH grant R35 GM139616). Data collection was partially supported by the Southeastern Center for Microscopy of MacroMolecular Machines (SECM4) (R24 GM145964).

## Data availability

Raw data for each relevant figure is available in 10.6084/m9.figshare.31460275.

## Author contributions

K.D.M. contributed to the protein and antibody expression and subsequent mAb characterization in vitro and in vivo. B.G.E. contributed to the cryo-EM analysis. M.A.E. contributed to the animal studies. A.L.M contributed to B cell sorting and mAb development. K.D.M. and J.J.M wrote the original manuscript. All authors reviewed and edited the final manuscript.

