## Supplemental Tables and Figures for "Molecular basis for protection and cross-protection by human antibodies targeting the parainfluenza virus hemagglutinin-neuraminidase protein"

| Table S1. Cryo-EM data collection, refinement, validation, and model-building statistics. | | | |
|---|---|---|---|
| | PIV3HN + PIV3HN-09 + 5217-2 | PIV3HN + PIV3HN-09 + PIV3HN-13 | PIV3HN + PIV3HN-13 + PIV3HN-05 |
| **Data collection and processing** | | | |
| Scope | Titan Krios | Titan Krios | Titan Krios |
| Magnification | 105,000 | 59,000 | 59,000 |
| Voltage (kV) | 300 | 300 | 300 |
| Electron exposure (e–/Å$^2$) | 60 | 60 | 60 |
| Defocus range (μm) | 0.8-2.8 | 0.8-2.8 | 0.8-2.8 |
| Camera | Gatan K3 | DE-Apollo | DE-Apollo |
| Pixel size (Å) | 0.88 | 0.79 | 0.79 |
| Symmetry imposed | C2 | C2 | C2 |
| Initial particle images (no.) | 3,800,000 | 150,000 | 1,200,000 |
| Final particle images (no.) | 238,000 | 22,000 | 287,000 |
| Map resolution (Å)<br>FSC threshold | 3.29<br>0.143 | 6.57<br>0.143 | 2.88<br>0.143 |
| Map resolution range (Å) | 2-6 | 5.5-10 | 2.3-7 |
| **Refinement** | | | |
| Initial model used (PDB code) | Alphafold3 | Alphafold3 | 7UR4/Alphafold |
| Model resolution (Å)<br>FSC threshold | 2.61<br>0.143 | 2.94<br>0.143 | 3.29<br>0.143 |
| Model resolution range (Å) | 2.8-6 | 2.4-6 | 2.5-8 |
| Map sharpening *B* factor (Å$^2$) | 116.1 | 84.0 | 135.1 |
| R.m.s. deviations<br>Bond lengths (Å)<br>Bond angles (°) | <br>0.003<br>0.564 | <br>0.003<br>0.588 | <br>0.003<br>0.717 |
| Validation<br>Clash-score<br>Poor rotamers (%) | <br>6.47<br>4.07 | <br>9.97<br>2.50 | <br>7.01<br>4.63 |
| Ramachandran plot<br>Favored (%)<br>Allowed (%)<br>Disallowed (%) | <br>90.99<br>8.56<br>0.45 | <br>95.04<br>4.96<br>0.00 | <br>86.82<br>12.61<br>0.56 |
| PDB accession code | TBD | TBD | TBD |
| EMDB accession code | TBD | TBD | TBD |
| EMPIAR accession code | TBD | TBD | TBD |

| Table S2. Nucleotide sequences of parainfluenza virus monoclonal antibodies. | |
|---|---|
| **5217-2 HC** | gaggtgcacctcctggagtctgggggaggcctggtccagcctgggggtcccgcagaatctcctgtgcagcctctggattctcttttggcagctatgccatgagctgggtccgccaggctccagggaaggggctggagtgggtctcgactataactagtagtggtgaaagcacagactacgcagactccgtgaagggccggttcatcatctccagagacaacgccaagaacacgctggatctcctaatgaacagcctgagagccggggacacggccgtttactactgtgcgaaggggggggcagtagtactgagcgctatggacgtctggggccaagggactacggtcaccgtctcctca |
| **5217-2 KC** | gacatccagttgacccagtctccatcttccctgtctgcatctgtaggagacagggtcaccatcacttgccaggcgagtcacgacattagtaactatctaaattggtatcagcagaaaccagggaaagcccctaagctcctgatctacgatgcatccaatttggaaacaggggtcccaccaaggttcagtggaggtggatctgggacacattttagtctcaccatcagcagcctgcagcctgaagacattgcaacatattactgtcaacaggatgatagtctcccattaactttcggccctgggaccaaggtggaaatcaaa |
| **5217-9 HC** | gaggtgcagctgttggagtcgggggaggcttggtacagcctgggggtccctgagactctcctgtgcagcctctggattcacctttagcagccatgccatgagctgggtccgccaggctccagggaaggggctggagtgggtctcaactattggtagtagtggcattagtacatactacacagactccgtgaagggccgcttcaccatctccagagacaattccaagaacacgctgtttcttcaactgaacagcctgagagccgaggacacggccgtctattactgtgaagtggggggggatcaaatcgcgtcttcctactggggccagggaaccctggtcaccgtctcctca |
| **5217-9 LC** | gacatccagatgacccagtctccttccaccctgtctgcatctgttggggatagagtcaccatcacttgccgggccagtgagaatattaatagttggttggcctggtatcagcagaaaccagggaaagcccctaaactcctgatctacaaggcgtctagtttacaaagtggggtcccatcaaggttcagcggcagtagatctgggacagaattcactctcaccatcagtagcctgcagcctgatgattttgcaacttattactgccaacagtataatacttatttctcgtacacttttggccaggggaccaagttggagatcaaa |

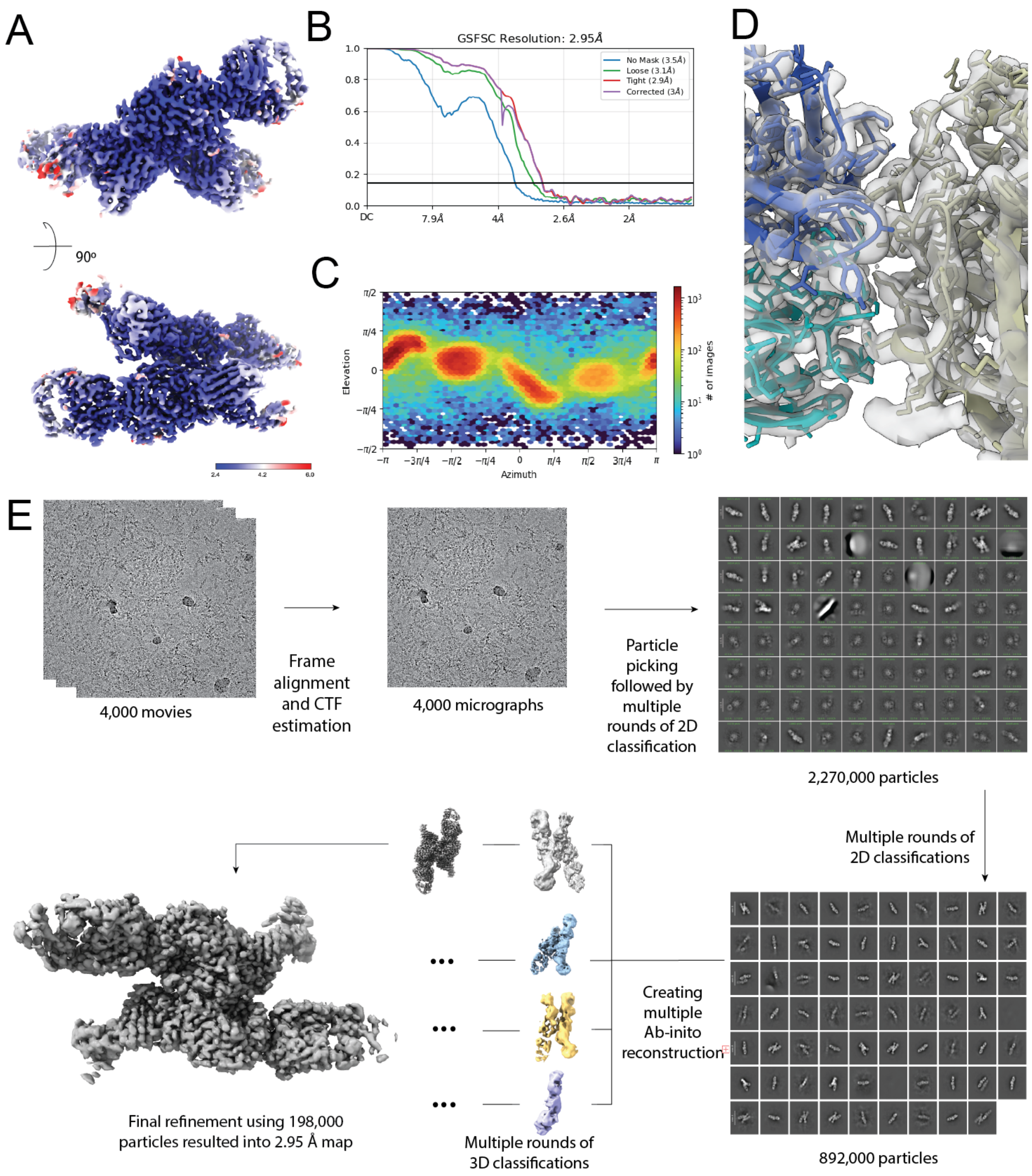


**Figure S2. PIV3HN-09 cryo-EM processing workflow.** (A) Local resolution map obtained for PIV3 HN protein bound to the PIV3HN-09 Fab and PIV3HN-13 Fab. (B) GSFSC curve of the refined cryo-EM map. (C) Particle distribution map for the final refinement. (D) Model to map fit example of the PIV3 HN protein interface with PIV3HN-09 Fab. (E) Overall cryo-EM data processing workflow.

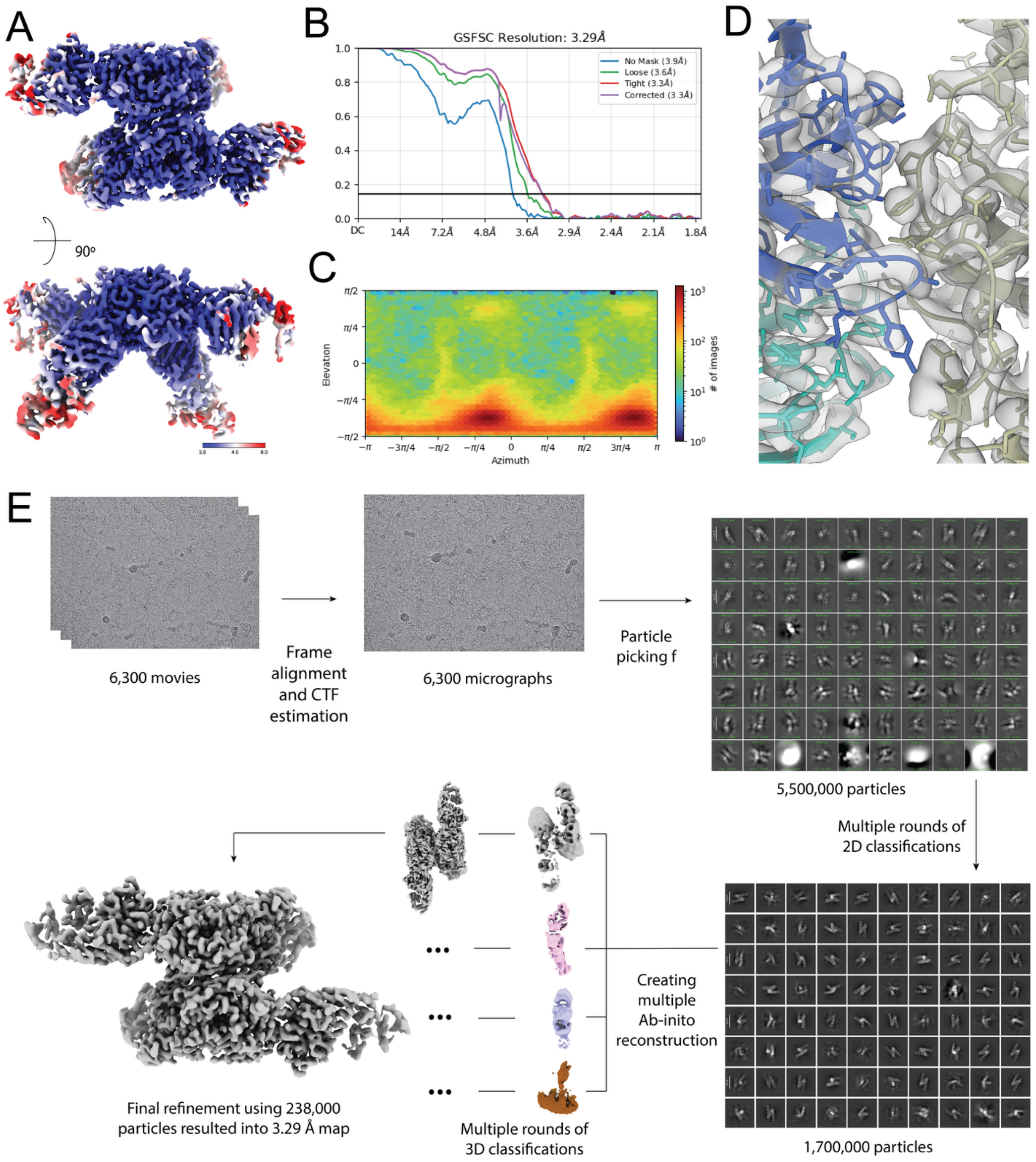


**Figure S3. 5217-2 cryo-EM processing workflow.** (A) Local resolution map obtained for PIV3 HN protein bound to 5217-2 and PIV3HN-09 Fabs. (B) GSFSC curve of the refined cryo-EM map. (C) Particle distribution map for the final refinement. (D) Model to map fit example of the PIV3 HN protein interface with 5217-2 Fab. (E) Overall cryo-EM data processing workflow.

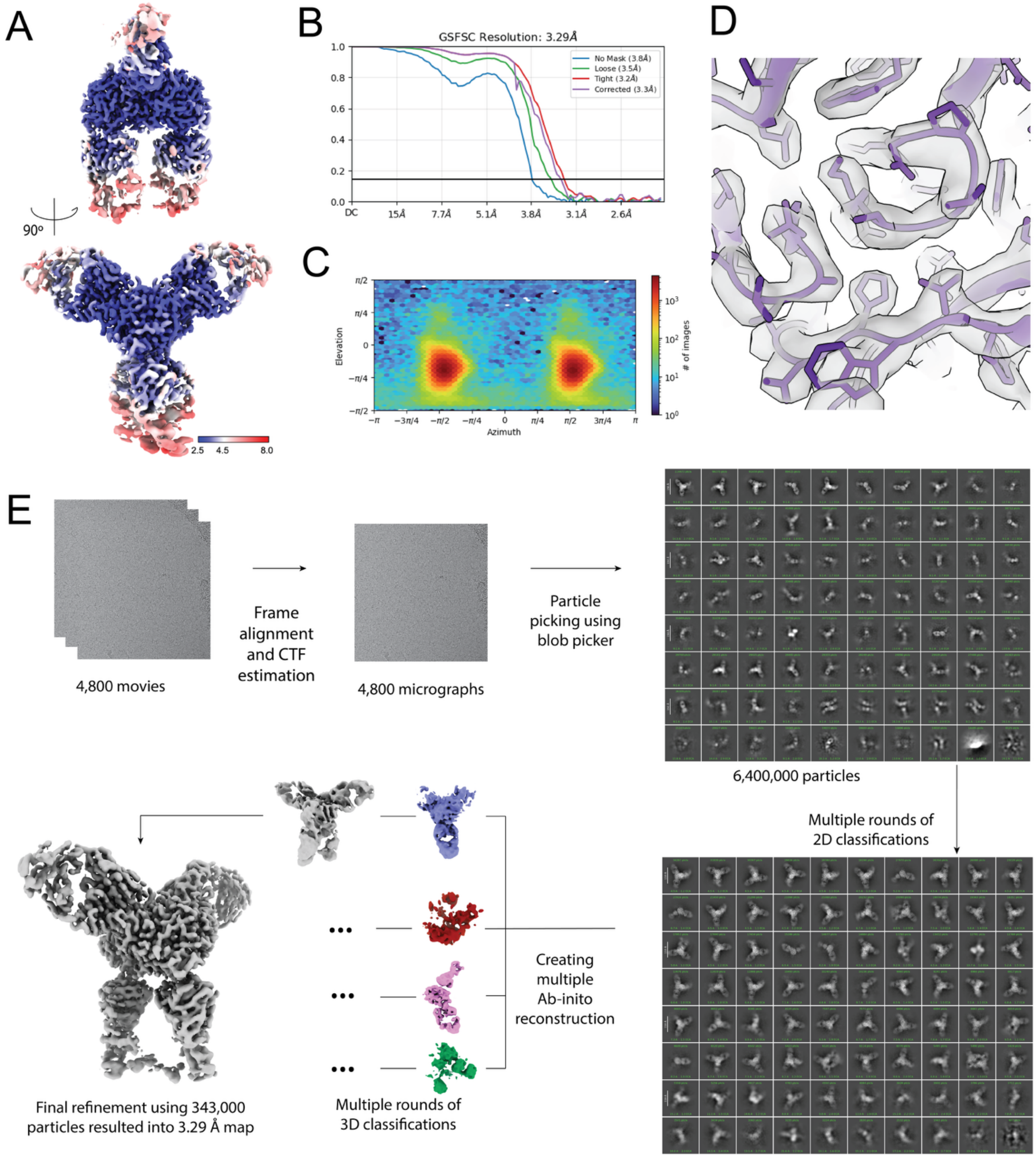


**Figure S4. PIV3HN-05 cryo-EM processing workflow.** (A) Local resolution map obtained for PIV3 HN protein bound to PIV3HN-05 and PIV3HN-13 Fabs. (B) GSFSC curve of the refined cryo-EM map. (C) Particle distribution map for the final refinement. (D) Model to map fit example of the PIV3 HN protein interface with PIV3HN-05 Fab. (E) Overall cryo-EM data processing workflow.
